# Genomic DNA repeats act as an antitoxin for a widespread Schlafen immune effector

**DOI:** 10.64898/2026.11.13.766530

**Authors:** Tori A. Boyle, Keegan Schoeller, Madeline R. Sanford, Nathan J. Hiestand, Catherine E. La, Hannah S. Aegerter, Barbaros Aslan, Asma Hatoum-Aslan

## Abstract

Many antiphage defenses employ toxic immune effectors in response to phage infection, necessitating robust strategies to counteract these toxins when cells are in the uninfected state. Here, we describe StaIN, the founding member of a unique family of toxin-mediated defenses that utilizes genomic DNA repeats for toxin restraint. The system consists of a repeat locus (*staI*) and a toxic protein (StaN) harboring Schlafen, ATPase, and helix-turn-helix (HTH) domains. We show that StaN is an ATP-activated tRNase that binds to *staI* DNA, and *staI* DNA is necessary to neutralize StaN’s toxicity. Computational analyses identified thousands of StaN homologs in bacteria and archaea that bear one or more HTH domains, and many reside in genomic loci that are enriched with DNA repeats. Moreover, functional analyses revealed that distant StaN homologs exhibit similar DNA-gated toxicity. Altogether, our findings expand the logic of toxin-antitoxin-like systems to include antitoxins composed of noncoding genomic DNA.

**In Brief:** DNA antitoxins store and restrain toxic Schlafen tRNases.

## Introduction

Bacteria have evolved a diverse array of immune systems to combat their viruses, phages. Some antiphage defenses protect individual infected cells by sensing and neutralizing the invading phage, whereas others protect the bacterial population by disrupting essential cellular components or pathways to halt phage replication^1^. The latter often involves the activation of defense-associated toxic proteins and results in cell growth arrest, dormancy, and/or death in a process known as abortive infection^2^. Indeed, toxic effectors are a recurring feature of many families of bacterial immune systems, thus necessitating robust molecular strategies to maintain these toxins in an inactive or restrained state in the absence of phage infection.

Toxin-antitoxin (TA) systems elegantly demonstrate the variety of mechanisms that bacteria employ to safely regulate potent toxic effectors. Typically consisting of a toxin that inhibits cellular growth and a cognate antitoxin that suppresses toxicity, TA systems play roles in plasmid maintenance, stress adaptation, persister cell formation, and antiphage defense^3–5^. These systems comprise four canonical types that are classified according to the molecular nature of the antitoxin and the mechanism by which it neutralizes the toxin: Types I and III TA systems encode RNA antitoxins that inhibit toxin expression or toxic protein function, whereas Types II and IV employ protein antitoxins that neutralize toxins either by direct binding or by indirectly counteracting toxin activity. Additional TA systems with RNA/protein antitoxins and distinct regulatory mechanisms (Types V-VIII) have expanded this framework^6–9^, and more recently, retrons have emerged as TA-like antiphage defenses that utilize reverse-transcriptases bound to DNA-RNA hybrids as antitoxins that neutralize a variety lethal enzymes^4,10^. While only a limited number of TA systems have been experimentally shown to provide antiphage defense, the advantage of harboring pre-made toxic effectors that are ready for deployment in response to phage infection suggests that TA systems dedicated to defense are more diverse and widespread than currently appreciated.

Here, we use genetic and biochemical approaches to identify and characterize StaIN, a unique TA-like antiphage system that relies on a cluster of DNA repeats (*staI*) to neutralize a toxic effector (StaN) bearing Schlafen (SLFN), Histidine Kinase ATPase (HATPase), and winged helix-turn-helix (wHTH) domains. We demonstrate that StaN possesses SLFN-mediated tRNase activity that requires ATP for allosteric activation, and binds to *staI* DNA through interactions with its wHTH domain. Computational analyses revealed that StaN homologs reside in diverse prokaryotes, bear one or more HTH domains, and are often encoded in genomic loci enriched with DNA repeats. Moreover, we show that distant homologs are functionally conserved, with toxicity that is neutralized by flanking DNA. Altogether, our findings define StaIN as the founding member of a widespread family of antiphage systems that store and restrain toxic Schlafen immune effectors on noncoding, often repetitive genomic DNA. More broadly, our findings expand the logic of toxin-antitoxin-mediated antiphage defense by showing that genomic DNA itself can function as an antitoxin.

## Results

### A repeat-associated defense locus in staphylococci protects against phage infection

With a longstanding interest in promoting the therapeutic use of phages to treat antibiotic-resistant infections, we continually seek to characterize yet unknown antiphage systems that may undermine the effectiveness of therapeutic phages targeting drug-resistant staphylococci. In light of our previous work demonstrating that antiphage defenses are concentrated in mobile genomic elements known as staphylococcal cassette chromosome (SCC) *mec* cassettes^11^, we screened for new defenses within the SCC*mec* regions of *S. epidermidis* clinical isolates^12^. Groups of SCC*mec*-encoded hypothetical genes were cloned into the high-copy plasmid pC194^13^, and constructs were introduced into two *Staphylococcus* backgrounds: *S. aureus* RN4220, a restriction-deficient strain^14^, and *S. epidermidis* LM1680, a “defenseless” variant of *S. epidermidis* RP62a that bears a large deletion encompassing the SCC*mec* cassette and all characterized antiphage defenses^15–17^. Notably, SCC*mec* elements are naturally capable of intra-genus transfer, making both backgrounds relevant proxies for the native strain. We then challenged the strains with diverse phages and identified a region in *S. epidermidis* 6593473 that protects against phages with myovirus morphology, particularly in the *S. aureus* background (Figure S1 A and B). These phages share ≥96% nucleotide identity with *Staphylococcus* phage K (Figure S2 A and B), a member of the *Twortvirinae* subfamily of the genus *Herelleviridae*^18^ that is commonly used in therapeutic applications^19^. The genomic region is also present in clinically-relevant *S. aureus* strains, including USA300. Thus, we sought to undertake more detailed characterization.

To pinpoint the gene(s) responsible for the defense phenotype, smaller subsets were introduced into *S. aureus* and the strains were re-tested. The results showed that a single predicted open reading frame (ACSVI7_RS00220) preceded by a cluster of eight 55-nucleotide tandem repeats are sufficient to reconstitute the defense phenotype (Figure 1A). Notably, the upstream repeated region bears homology to *tsr1*, a small noncoding RNA with unknown function identified in *S. aureus*^20^, and multiple attempts to clone the open reading frame in the absence of repeats proved unsuccessful. Thus, we speculated that that the defense-associated gene may be toxic on its own and works together with the repeats to mediate defense.

**Figure 1.**
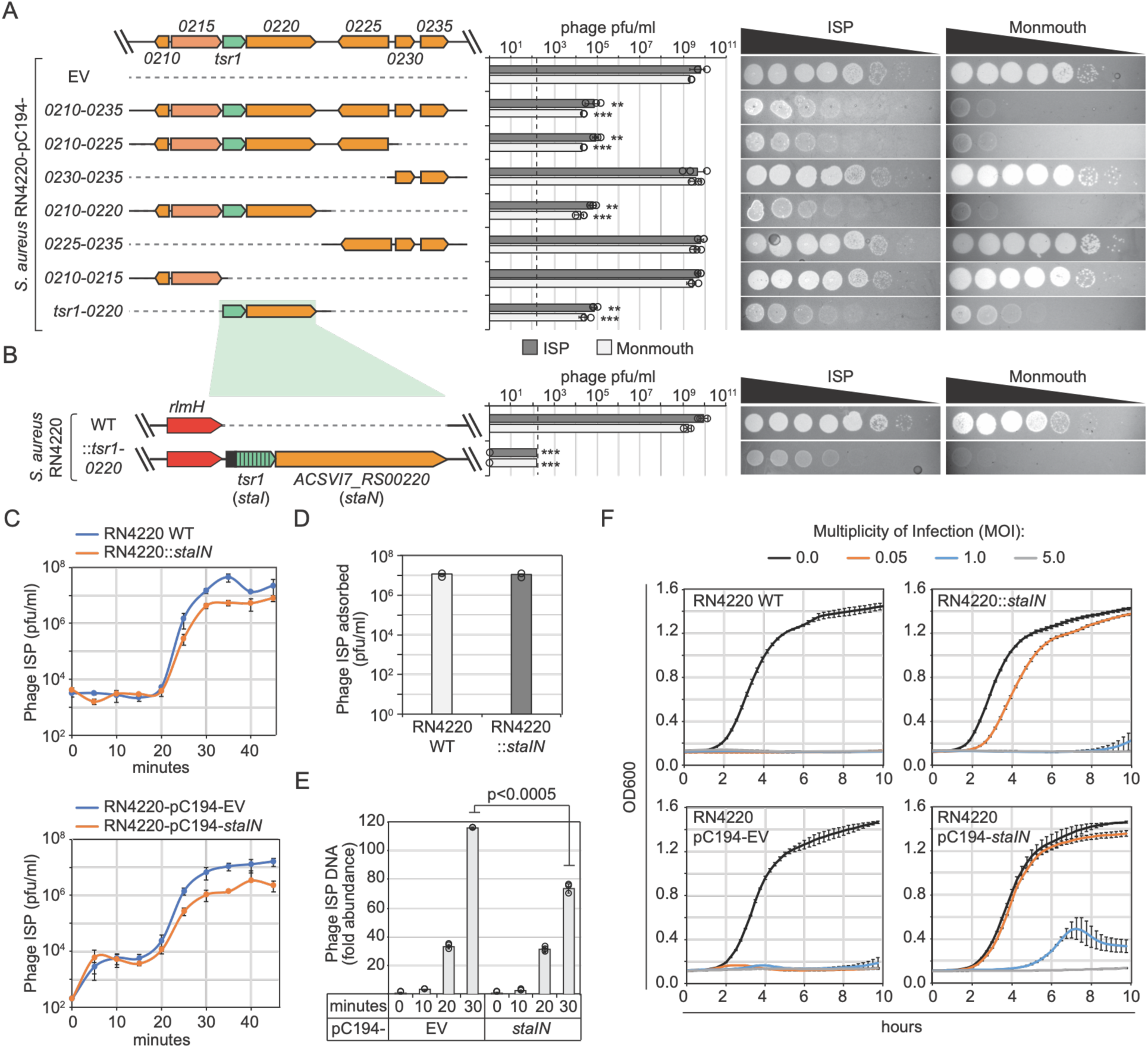
A repeat-associated defense locus in staphylococci (*staIN*) protects against K-like myophages. **(A)** Segments of the *S. epidermidis* 6593473 genome responsible for antiphage defense activity and corresponding plasmids (pC194-) containing them. Cells bearing indicated plasmids were challenged with 10-fold dilutions of phages, and average plaque-forming units per milliliter (pfu/ml) from three technical replicates is shown (+/-SD) with representative plate images. Dotted line, limit of detection. Asterisks indicate p values < 0.01 (**) and < 0.001 (***) from two-tailed Student’s t-tests. **(B)** The minimum genomic locus responsible for defense activity was introduced into *S. aureus* RN4220 downstream of *rlmH* and challenged with indicated phages as in panel A. **(C)** One-step growth curves with ISP following infection of strains carrying *staIN* in a single genomic copy (top) or plasmid (bottom). **(D)** Adsorption assay following ISP challenge of *S. aureus* RN4220::*staIN.* **(E)** Quantitative PCR (qPCR) measurements showing relative abundance of ISP DNA at different time points following infection. Measurements from three technical replicates are shown (+/-SD). **(F)** Bacterial growth dynamics following ISP infection at different multiplicities of infection (MOI, phage:bacteria ratio). OD600 (optical density at 600 nM) measurements from three technical replicates are shown (+/-SD).

To test defense function in a more physiologically-relevant context, we introduced the *tsr1*-ACSVI7_RS00220 locus into *S. aureus* in a single copy downstream of *rlmH*, where SCC*mec* cassettes naturally integrate^11^, and repeated the phage challenge assays. Unexpectedly, a more robust defense phenotype was observed, with complete elimination of plaques (Figure 1B). From these data, we concluded that *tsr1*-ACSVI7_RS00220 constitutes an antiphage defense system, and based on its architecture and presumed toxicity, we posited that the system may function similarly to Type III toxin-antitoxin systems, in which RNA antitoxins, often composed of repetitive sequences, bind and inhibit toxic protein(s) encoded downstream^21,22^. Accordingly, we named the system *staIN*, consistent with the proposed naming convention for Type III TA systems, wherein *staI* refers to the <u>sta</u>phylococcal inhibitory genetic locus, and *staN* refers to the gene that encodes the <u>sta</u>phylococcal toxi<u>n</u>.

We next performed a series of experiments to determine the impact of *staIN* on the growth dynamics of ISP and its *S. aureus* host. First, one-step growth curves showed that *staIN* significantly reduces phage amplification by 11-17 – fold (Figure 1C). Second, adsorption assays revealed that the system has no effect on phage attachment to cells (Figure 1D). Third, quantitative PCR (qPCR) showed that phage DNA entry occurs in the presence of *staIN*; however, the system causes is a significant (∼40%) reduction in phage DNA accumulation by ∼30 minutes post-infection (Figure 1E). Finally, phage challenge assays in liquid culture with different phage:bacteria ratios (i.e. multiplicities of infection (MOIs)) showed that *staIN* supports bacterial growth only at a low MOI (<1) (Figure 1F). From these data, we concluded that *staIN* is deployed after phage DNA injection and provides population-level defense (i.e. abortive infection).

### staN encodes a toxin, and staI DNA, but not RNA, neutralizes staN’s toxicity

Our inability to clone *staN* under its native promoter in the absence of *staI* repeats hinted that *staN* is toxic, and the repeats might be necessary to mitigate toxicity. To test this directly, we inserted *staN* in the presence/absence of repeat(s) into plasmid pTET^17^ downstream of an anhydrotetracycline-(aTc-) inducible promoter. Dilutions of cells harboring the plasmids were then plated +/- aTc, and colony-forming units per mililiter (cfu/ml) were enumerated the following day. The results showed a ∼1,000-fold reduction of colony counts in cells expressing *staN* alone, indicating the gene is indeed toxic; however, in the presence of the upstream repeats, toxicity is completely eradicated (Figure 2A), consistent with toxin-antitoxin-like activity. We also conducted this assay with constructs bearing progressively fewer repeats and observed that a minimum of two repeats are necessary for full toxin neutralization (Figure 2A).

**Figure 2.**
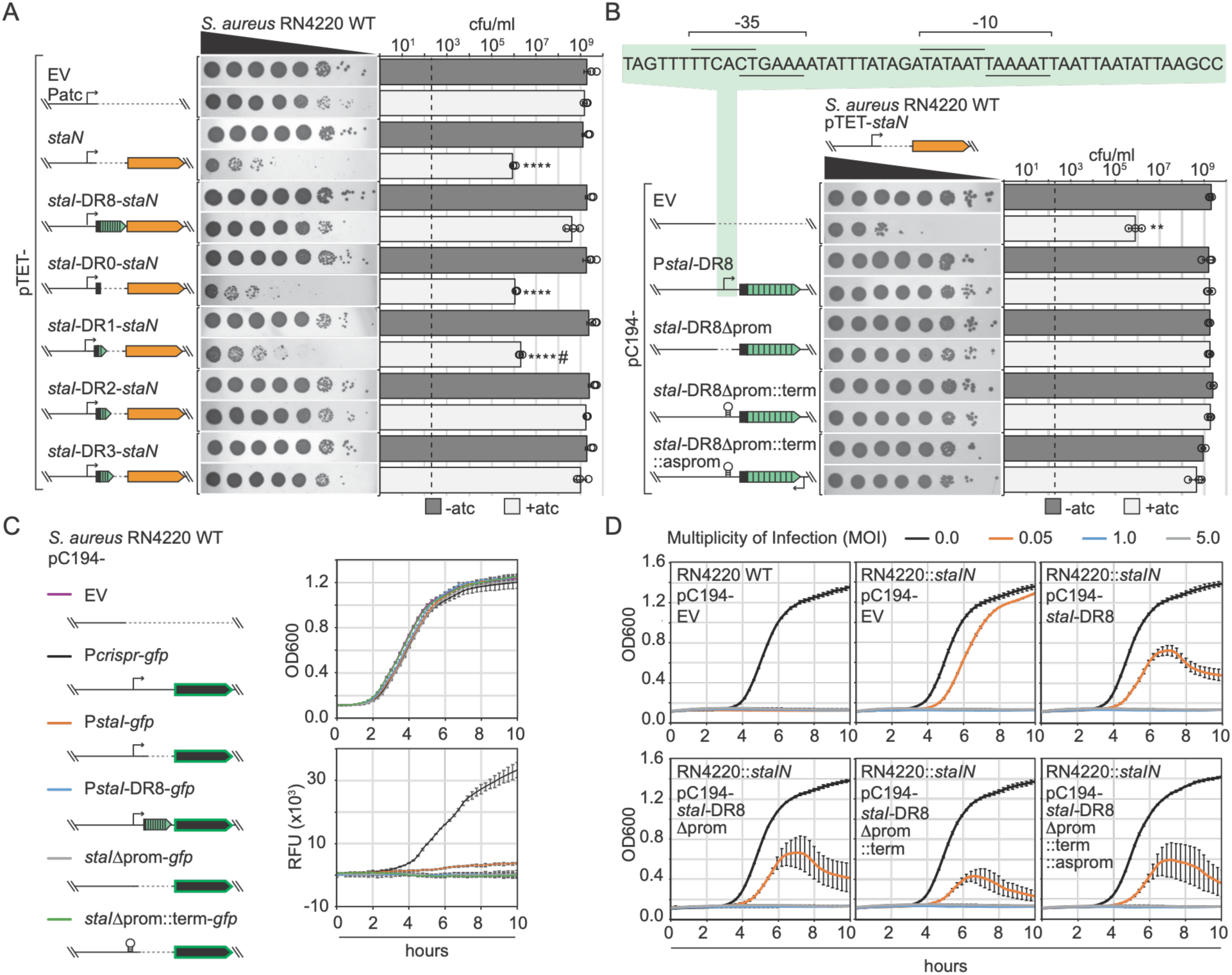
StaN is toxic to cells, and *staI* DNA (not RNA) mitigates toxicity. **(A)** Toxicity assays with *staN*. 10-fold dilutions of indicated strains were plated +/-anhydrotetracycline (aTc). Average colony-forming units per milliliter (cfu/ml) from three technical replicates (+/-SD) and corresponding plate pictures are shown. #, tiny colonies observed out to the 10^-7^ dilution. Dotted line, limit of detection. Asterisks indicate p values < 0.0001(****) from two-tailed Student’s t-tests. **(B)** Dual-plasmid toxicity assays with *staN* and s*taI* variants cloned into separate plasmids were performed as in panel A. Asterisks indicate p values < 0.001(**) from two-tailed Student’s t-tests. The *staIN* promoter region is displayed above the graph with-10 and-35 elements delimited by lines. **(C)** Cell growth and GFP reporter assays. Indicated strains were grown in liquid culture, and their growth (optical density at 600 nm, OD600) and GFP reporter activity (relative fluorescence units, RFUs) were measured every 15 minutes. Average readings from three technical replicates are shown (+/-SD). **(D)** Cell growth dynamics assays with ISP infection. Average OD600 measurements from three technical replicates are shown (+/-SD) as a representative of three independent trials.

We next sought to determine whether the antitoxin functions as DNA or RNA. We developed a dual-plasmid system in which the *staI* locus—including two predicted promoters, a short leader and all eight repeats—was inserted into pC194, and the construct was introduced into *S. aureus* harboring pTET-*staN* (Figure 2B). Cells containing both plasmids were then plated with aTc to confirm that the *staI* locus maintains antitoxin activity when supplied *in trans*. Next, we introduced a series of mutations in the *staI* locus that maintain the leader and repeat sequences, but progressively reduce/eliminate transcript production. These constructs harbor i) a deletion of the *staI* promoter(s), ii) an insertion of a terminator to replace the promoter(s), and/or iii) an insertion of an antisense promoter downstream of the repeats to generate antisense transcripts that would bind any remaining transcripts produced in the forward direction. These constructs were introduced into *S. aureus* harboring pTET-*staN* and the toxicity assay was repeated. Remarkably, each of these constructs was similarly able to restore cell growth under inducing conditions (Figure 2B), indicating that *staI* DNA, but not RNA, provides antitoxin activity.

With DNA serving as an antitoxin, *staIN* falls outside the current classification of canonical TA systems, which rely on antitoxins composed of proteins or RNAs^4^. Nonetheless, we sought to explore other parallels to Type III TA systems as the closest functional representatives. Type III TA systems typically encode intrinsic terminators downstream of the repeats to prevent excess toxin production^23^; however, we were unable to locate such a terminator between *staI* and *staN* using *in silico* analyses^24^. Thus, to qualitatively assess expression levels, we created green fluorescent protein (*gfp*) fusions of the *staI* promoter (P*staI-gfp*) or the promoter followed by eight repeats (P*staI*-DR8-*gfp*) (Figure 2C). As negative controls, reporters with the *staI* promoter deleted or a terminator inserted in place of the promoter were constructed; and as a positive control, *gfp* was fused downstream of the constitutively active promoter that drives expression of the Type III-A CRISPR-Cas system in *S. epidermidis* RP62a^25,26^. The reporter strains were grown in liquid culture, and measurements of cell growth and promoter strength (relative fluorescence units (RFUs)) were taken periodically over a 10-hour period. As expected, the negative controls showed no fluorescence output, further validating that the antitoxin must be DNA. Moreover, we observed a modest fluorescence output for P*staI-gfp*, and no detectible fluorescence from P*staI*-DR8-*gfp* (Figure 2C), suggesting that *staIN* expression levels are low under standard growth conditions, and transcription may be further hindered by the repeats.

TA systems must maintain a delicate balance of toxin and antitoxin levels to restrain toxin activity, yet ensure robust toxin activation under antitoxin-depleting conditions. In agreement, we observed that a single *staI* repeat is insufficient to mitigate *staN* toxicity (Figure 2A), and on the opposite end of the spectrum, we wondered whether an overabundance of repeats would overwhelm the system with antitoxin and interfere with the system’s function. To test this, constructs containing WT or mutant versions of the *staI* locus with the presence/absence of functional promoter(s) (Figure 2B) were introduced into *S. aureus* RN4220::*staIN*. Each construct harbors eight direct repeats, and the plasmid backbone (pC194) has an estimated copy number of 15^13^, thus providing ∼120 additional *staI* repeats *in trans*. These strains were then challenged with ISP at varying MOIs in liquid culture (Figure 2D), and the results showed that all constructs indeed inhibit antiphage defense to a similar degree. Altogether, our findings demonstrate that while *staIN* does not abide by the composition of canonical TA systems, it employs toxin-antitoxin-like logic to defend against phages, with the antitoxin composed of noncoding genomic DNA.

### StaN is a Schlafen domain-containing effector with ATP-dependent tRNase and DNA-binding activities

We next explored the biochemical basis of StaN’s toxicity and its relationship with *staI* DNA. First, InterProScan^27^ was used to identify three domains: an N-terminal SLFN_AlbA_2 domain (SLFN) predicted to have tRNase activity^28,29^, a Histidine Kinase ATPase domain (HATPase), and a C-terminal winged helix-turn-helix (wHTH) DNA binding domain (Figure 3A). The search also indicated that StaN belongs to the GLPR-related transcriptional repressor family (defined in the PANTHER database^30^). Second, we aligned the amino acid sequences of hundreds of StaN homologs (Table S1) and generated a histogram to reveal conserved regions. These analyses identified highly-conserved amino acids that map to the predicted catalytic domains (Figure 3A). Finally, AlphaFold 3^31^ (AF3) was used to generate a structural prediction of StaN bound to two Mg^2+^ ions and ATP. Consistent with predicted nuclease and ATPase activities, the results returned a high-confidence structure, with one Mg^2+^ ion bound in the SLFN domain, and Mg^2+^ and ATP bound to the HATPase domain (Figure 3B). In light of these predictions, we speculated that StaN’s SLFN domain may catalyze tRNA degradation, ATP may be needed for this activity, and the wHTH domain may facilitate binding to *staI* DNA.

**Figure 3.**
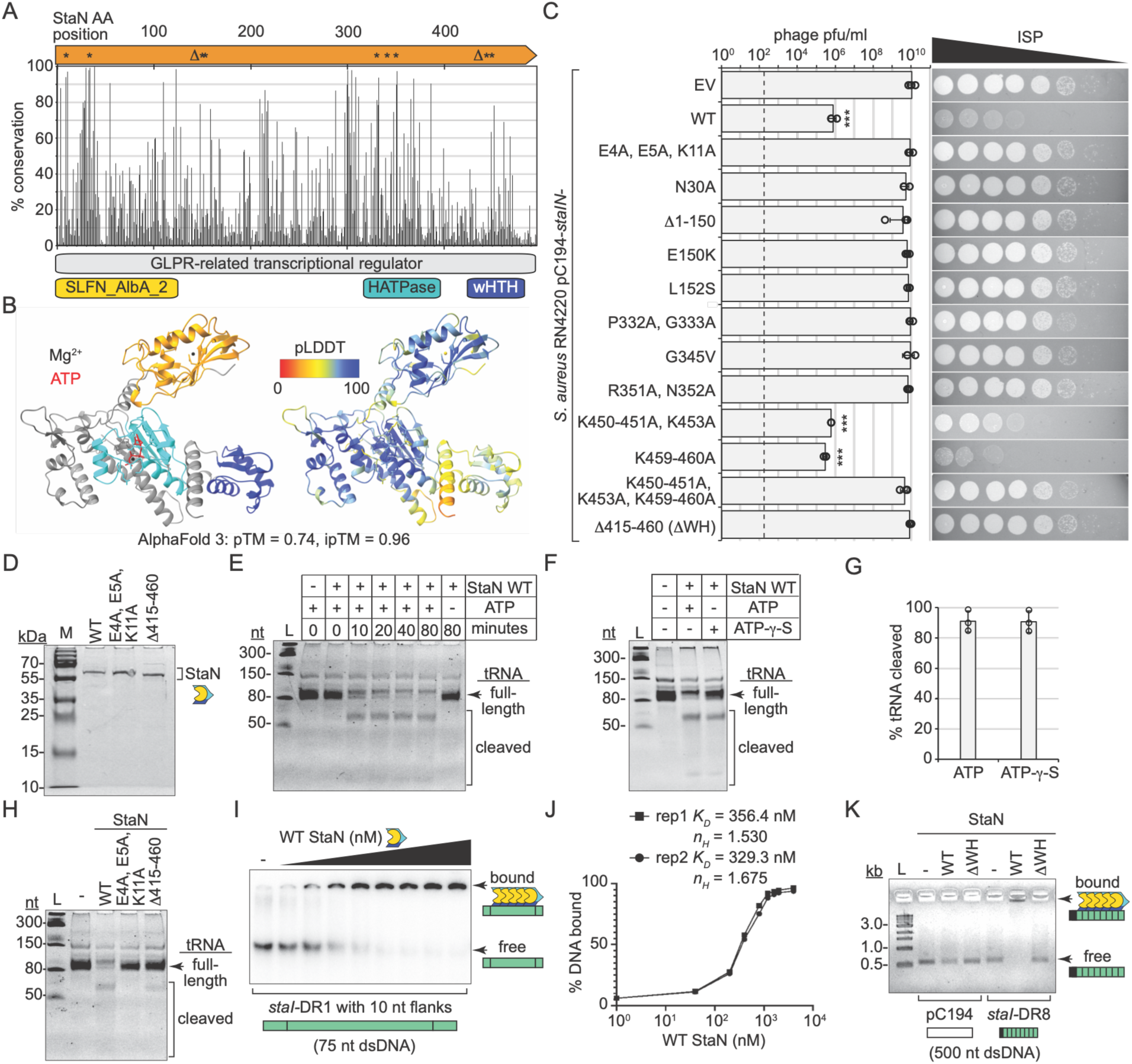
StaN exhibits ATP-activated tRNase activity and binds to *staI* DNA. **(A)** Fraction of 717 StaN homologs that possess amino acids (AA) identical to StaN at each position. Domains predicted by InterProScan (bottom) and positions where point mutations (*) or deletions (Δ) were introduced (top) are shown. **(B)** AF3 models of StaN bound to ATP and 2 Mg^2+^ ions, colored according to predicted domains (left) and Local Distance Difference Test (pLDDT) scores (right). **(C)** Phage challenge assays with cells bearing WT *staIN* and indicated mutants. Average pfu/ml of three technical replicates are shown (+/-SD) with representative plate pictures. Dotted line, limit of detection. Asterisks indicate p values < 0.001 (***) from two-tailed Student’s t-tests. **(D)** Purified WT and mutant StaN variants resolved on an SDS-PAGE gel. **(E)** tRNase assays in which *E. coli* tRNAs were incubated with WT StaN +/- ATP for indicated times and resolved on a Urea-PAGE gel. Representative of three independent trials is shown. **(F)** tRNase assay in which WT StaN was incubated with ATP or ATPγS for 40 minutes. **(G)** Average fraction tRNAs cleaved (+/- SD) from three independent trials of the experiment shown in panel F. **(H)** tRNase assays with indicated StaN variants in the presence of ATP. **(I)** DNA-binding assays with 75-nt *staI* repeat double-stranded DNA (dsDNA). DNA was mixed with increasing concentrations of WT StaN for 15 minutes and resolved on a Native-PAGE gel. **(J)** Quantification of the experiment in panel I. Two independent trials were plotted, and dissociation constants (K_D_) and Hill Coefficients (n_H_) were calculated for each trial. **(K)** DNA binding assays in which indicated dsDNA probes were mixed with WT StaN or the ΔWH variant for 15 minutes and resolved on an agarose gel.

To begin to test this, we introduced mutations in conserved domains in the pC194-*staIN* construct (Figure 3 A and C). We included three spontaneous mutants that arose during initial attempts to clone *staN* in the absence of repeats, some of which occur outside of these domains (E150K, L152S, and G345V). Deletions of the entire N-terminal SLFN or C-terminal wHTH domain were also created, and all constructs were tested for antiphage activity. We observed that the majority of mutants failed to protect against phage infection (Figure 3C), suggesting that all domains are necessary for function. However, we cannot rule out the possibility that loss of defense may be attributed to reduced gene expression from the mutant constructs.

Thus, we next sought to test StaN’s biochemical functions more directly. We overexpressed and purified WT StaN and two variants harboring mutations in the SLFN (E4A, E5A, K11A) and wHTH domains (Δ415-460 or ΔWH) using a four-step process that includes a final size exclusion chromatography (SEC) step (Figure S3A). The SEC elution profile revealed that StaN purifies in a monomeric state, with trace levels of dimers and higher-order oligomer(s) (Figure S3B). We also noted the presence of smaller protein species that were confirmed to constitute StaN degradation products by mass spectrometry. The E4A, E5A, K11A mutant exhibited an elution profile similar to that of WT (Figure S3C), while the ΔWH did not produce a dimer peak, suggesting that the wHTH domain may be necessary for oligomerization (Figure S3D).

A series of biochemical assays were next performed to probe StaN’s activities using concentrated SEC fractions containing purified full-length proteins (Figure 3D). First, WT StaN was incubated with Mg^2+^, ATP, and a mixture of tRNAs to test for tRNase activity. We observed that StaN can indeed cleave tRNAs, but only in the presence ATP (Figures 3E and S4). This was surprising considering previously-reported prokaryotic SLFN domain-containing proteins (pSlfns), which exhibit tRNase activity in the absence of ATP^28,29^. Thus, we speculated that rather than driving catalytic function, ATP is likely required for StaN’s allosteric activation. To test this, tRNase assays were repeated with ATPγS (adenosine 5’-O-(3-thiotriphosphate)), an analogue that is weakly hydrolyzable. The results showed that ATPγS was able to support tRNase activity to the same degree as ATP (Figure 3 F and G), indicating that ATP likely functions allosterically. Moreover, tRNase activity was completely absent and significantly diminished in the E4A, E5A, K11A and ΔWH mutants, respectively (Figure 3H). From these data, we concluded that StaN’s SLFN domain has ATP-activated tRNase activity, and an intact wHTH domain is necessary to support its full function.

Finally, we tested for DNA binding using gel shift assays. Increasing amounts of WT StaN were first combined with a radiolabeled double-stranded DNA (dsDNA) probe comprising a single *staI* repeat with 10 nucleotide flanks. After a brief incubation period, the mixtures were resolved on native polyacrylamide gels. The results showed a progressive upward shift of the probe in the presence of increasing concentrations of StaN, characteristic of a DNA-binding protein (Figure 3I). Moreover, binding curves indicated a disassociation constant in the nanomolar range, and a Hill coefficient > 1, indicating cooperativity (Figure 3J). To more closely simulate the native genomic locus and investigate sequence specificity, gel shift assays were performed with 500-nucleotide dsDNA PCR amplicons that encompass either the *staI* locus or a nonspecific segment of pC194. The results showed a robust upward shift of the *staI* repeat DNA, and not the non-specific DNA in the presence of WT StaN (Figure 3K). Moreover, a shift was not observed following incubation with the ΔWH mutant. From these observations, we concluded that StaN binds specifically to *staI* repeat DNA through interactions with its wHTH domain.

### A phage-encoded putative DNA binding protein promotes staN-mediated toxicity

We next wondered how StaIN becomes activated upon phage infection. Providing potential clues, the well-characterized T4-like myophages of *E. coli* are known to degrade host genomic DNA in the early stages of infection, and one study showed that *Staphylococcus* phage K, a closer relative of ISP (Figure S2), may employ a similar strategy to liberate free nucleotides for incorporation into newly-synthesized phage^32^. However, the nucleases involved in host DNA degradation by phage K have yet to be experimentally verified^33^. Thus, we sought to preliminarily test for ISP-mediated DNA degradation by monitoring *S. aureus* DNA integrity following challenge with the phage. The results showed no evidence of DNA degradation by phage ISP at up to 20 minutes post-infection (Figure S5A). In contrast, the same assay performed with *E. coli* phage T4 clearly showed DNA fragmentation starting 5-10 minutes post-infection (Figure S5B). Thus, we speculate that global host DNA degradation is unlikely to play a role in StaN deployment following ISP infection.

To explore alternative possibilities, we next sequenced ISP “escapers” with spontaneously acquired mutation(s) that allow phages to overcome immunity. Twelve such escapers were purified and their genomes were subjected to Illumina sequencing. Illumina reads were then aligned to a *de novo* assembly of the ancestral phage genome (Figures 4A and S6), revealing that all escapers harbor mutation(s) in a common genetic locus encoding gene product 85 (*gp85*), a hypothetical protein (Figure 4B). ISP Gp85 is highly-conserved among phages belonging to the *Herelleviridae* Family, and the equivalent protein in Phage K, Gp102, is expressed immediately after infection and thought to be involved in host takeover or lysis timing^34,35^. Moreover, although its exact function is unknown, the protein is hypothesized to work through DNA binding. In agreement, an InterProScan search for conserved domains in Gp85 identified an N-terminal homeodomain-like domain, similar to homeobox domains found in DNA-binding proteins and transcription regulators in vertebrates^36^. Also, we used AF3 to model complexes of Gp85 bound to sequences encompassing 60 predicted promoters within ISP, and many complexes exceeded the thresholds for confident structures and interactions (Table S2). These structures displayed the homeodomain-like domain in direct contact with the major groove of the DNA (Figure S7 A and B), consistent with sequence-specific DNA binding.

**Figure 4.**
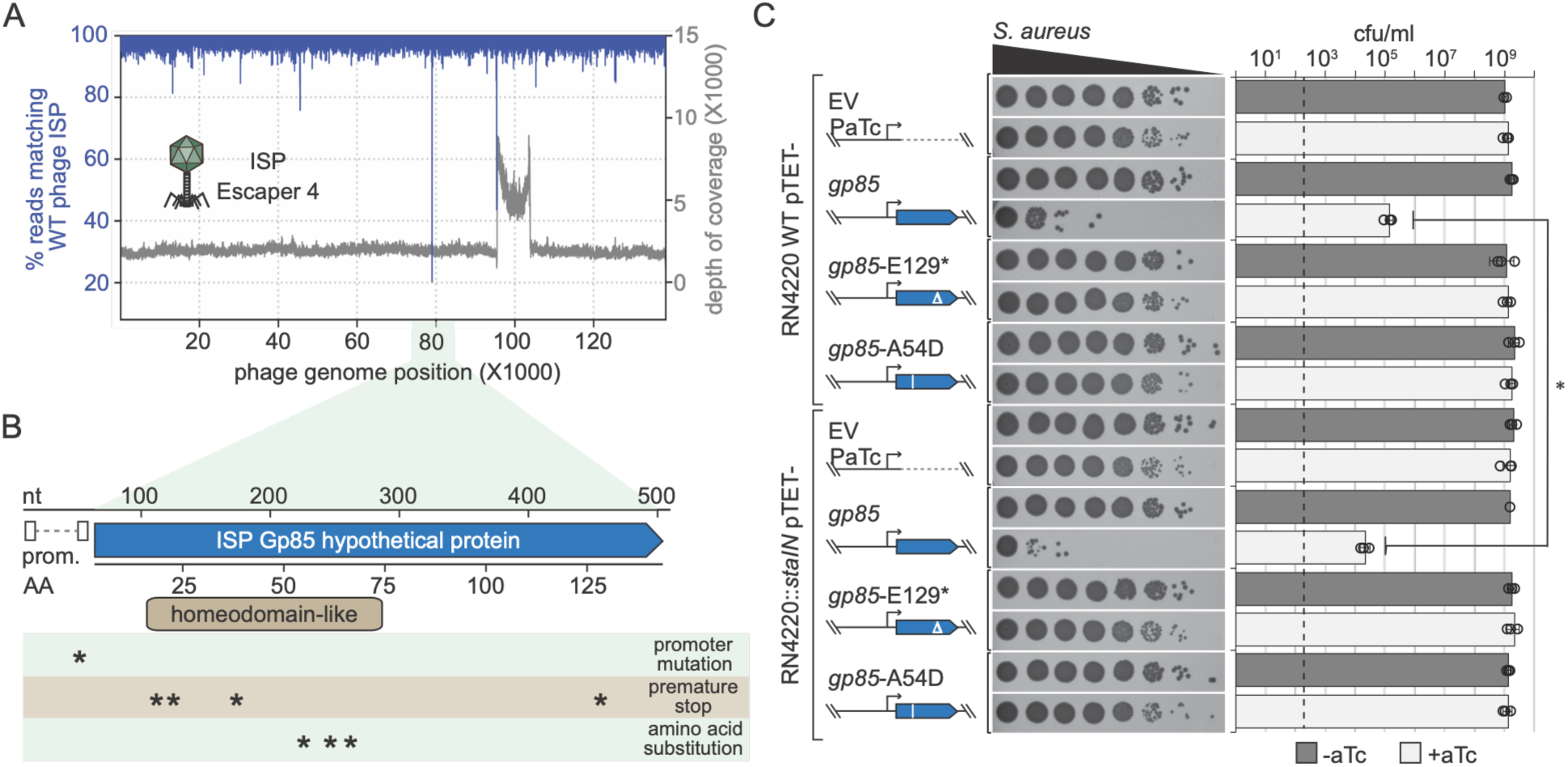
A phage-encoded transcription regulator promotes StaIN toxicity. **(A)** Illumina sequencing reads from one representative ISP escaper aligned with the *de novo* assembly of ancestral ISP. **(B)** Summary of mutations in ISP Gp_85 across 12 escaper phages. Asterisks show positions of indicated mutations. The homeodomain-like domain is highlighted. **(C)** Cell toxicity assays with *gp85*. 10-fold dilutions of cells bearing the indicated plasmids were plated +/- anhydrotetracycline (aTc). Average colony-forming units per milliliter (cfu/ml) from three technical replicates (+/- SD) and corresponding plate pictures are shown. Dotted line, limit of detection. Asterisks (*) indicate p value < 0.05 determined by a two-tailed Student’s t-test.

To test whether ISP *gp85* alone induces *staIN*-mediated toxicity, we inserted the coding regions for WT Gp85 and two escaper phage variants (E129* and A54D) into pTET, and the constructs were introduced into *S. aureus* RN4220 WT or RN4220::*staIN*. These strains were then grown in the presence of aTc, and the results showed that while induction of both mutant versions of *gp85* have no effect on cell growth, WT *gp85* in the absence of *staIN* caused a significant ∼10,000-fold reduction in colony counts (Figure 4C), indicating that Gp85 alone is toxic. This observation is consistent with a previous report on Phage K Gp102 and its connection with a temperature-sensitive phage growth phenotype^34^. However, despite the background toxicity caused by Gp85 alone, when the protein is expressed in RN4220::*staIN*, colony counts are further significantly reduced ∼10-fold, indicating that *gp85* expression promotes *staIN*-mediated toxicity in the absence of any other phage components. These data suggest that Gp85 may be sufficient to activate StaIN defense.

We next sought to purify Gp85 and test for direct interactions with StaN; however, multiple attempts proved unsuccessful, likely owing to Gp85’s toxicity. In lieu of this, we performed a series of *in silico* analyses using AF3, and found that while Gp85 and StaN are unlikely to interact with each other, both exhibit high-confidence interactions with one or more segments of the *staI* locus (Table S3). Consistent with experimental observations (Figure 3 I and J), many of the models with StaN bound to *staI* DNA showed StaN’s wHTH domain contacting the DNA (Figure S7C), and one confident structure with Gp85’s homeodomain-like domain bound to *staI* DNA was found, interestingly, on the opposite face (Figure S7D). These results hint at the compelling possibility that Gp85 may bind and displace StaN at the *staI* locus; however a definitive determination will require further experimentation.

### StaN homologs are widespread, possess one or more HTH domains, and are encoded in proximity to genomic loci enriched with DNA repeats

We next developed a computational pipeline to investigate the phylogeny and domain architectures of StaN homologs and the extent to which they are encoded proximal to DNA repeats (Figure 5A). First, BLASTp was used to identify ∼5000 StaN homologs in the RefSeq protein database (Table S4). Homologs were clustered using MMseqs2, yielding 2,293 cluster representatives. Each representative was folded using AF3, and resulting structural domains were parsed and classified using DPAM^37^ (Domain Parser for AlphaFold Models) and the ECOD (Evolutionary Classification of Protein Domains) database^38^, respectively. Since cluster members were 98.4% (median) identical to their representatives, each cluster member was assigned the domain architecture of its representative. After filtering out truncated variants, 4,036 intact homologs in 1,668 clusters remained.

**Figure 5.**
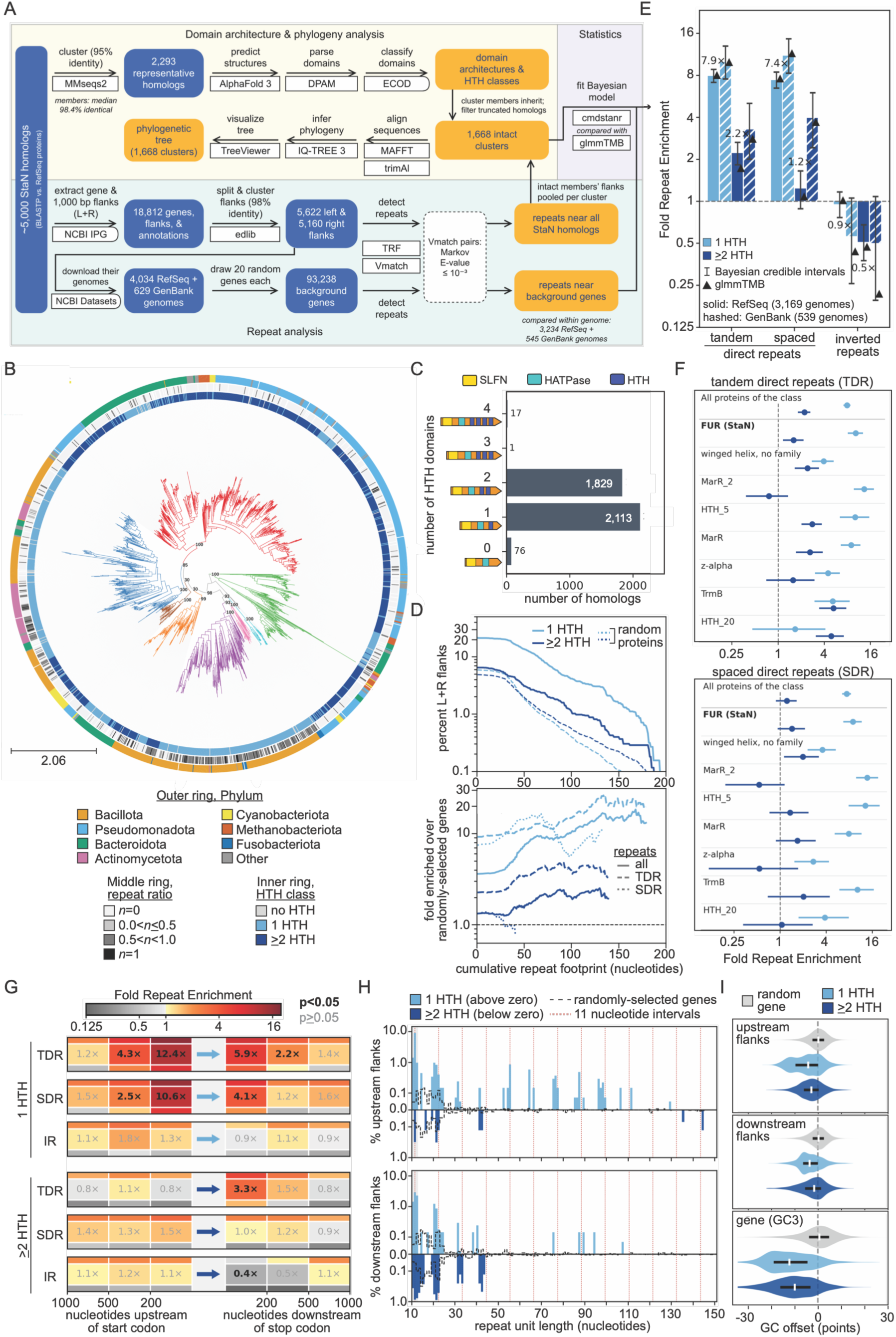
Phylogenetic distribution, architecture, and repeat enrichment across StaIN systems. **(A)** Schematic of the computational analysis workflow (see methods). **(B)** Phylogenetic tree of StaN homologs. Bootstrap values are shown for major branches. Phylum of cluster representative (outer ring), fraction of cluster members with repeats within 200 nucleotides (middle ring), and number of HTH domains in each homolog (inner ring) are shown. **(C)** Distribution of StaN homologs with indicated numbers of HTH domains. **(D)** Fraction of flanks of StaN homologs and randomly-selected (background) proteins with at least a given repeat-covered length within 200 nucleotides. Homologs and background genes (top) and their ratios (bottom) are shown. **(E)** Fold enrichment of flanks with a repeat within 200 nucleotides of the homolog over background genes. Bars show posterior medians with 95% credible intervals; triangles show glmmTMB point estimates. Genome counts include only genomes with an HTH+ or HTH++ flank searched across the full 200 nucleotides. **(F)** Fold enrichment by ECOD HTH family, for families with ≥20 clusters in ≥10 genera in both HTH classes. **(G)** Fold enrichment by side and distance from homologs. Outer stripes are 95% bootstrap bounds. **(H)** Share of flanks carrying a tandem array of indicated repeat unit length upstream (top) and downstream (bottom) of homologs. **(I)** GC content of the upstream and downstream flanks, and the gene’s third codon positions (GC3), relative to background genes of the same genome.

To infer phylogeny, sequences of cluster representatives were aligned using MAFFT^39^, alignments were trimmed with trimAl^40^, and a phylogenetic tree was built using IQ-TREE 3^41^. These analyses identified homologs in bacteria and archaea (Table S4), with the highest representation in bacterial phyla Pseudomonadota and Bacillota, and archaeal homologs in Methanobacteriota (Figure 5B, outer ring). We observed that the majority of intact StaN homologs (52%) carry a single C-terminal HTH domain (hereafter HTH+), and remarkably, a large fraction (46%) carry two or more (hereafter HTH++), almost always of different ECOD families (Figure 5B, inner ring, and C). Notably, a similar domain analysis using InterProScan on the same representatives failed to detect multiple HTH domains in the majority of HTH++ representatives, even though most of HTH ECOD families have corresponding protein families defined in the Pfam database (which InterProScan searches). Nonetheless, we spot-checked the AF3 structures of a subset of HTH++ representatives to confirm they indeed possess multiple HTH domains.

We next searched genomic contexts of each homolog for the presence of DNA repeats. We extracted up to 100 genomic contexts of each homolog (coding region plus 1000 nucleotides on both sides) available through the NCBI Identical Protein Groups (IPG) database, yielding 18,812 genome segments (Table S5). The segments were split into coding regions and flanks, and near-identical flanks for each homolog were merged. Flanks were then searched for the presence of direct and inverted repeats using i) Tandem Repeats Finder^42^ (TRF) to detect direct tandem repeats, and ii) Vmatch^43^ to detect direct spaced and inverted spaced repeats (i.e. repeats with up to 300 nucleotides in between them). In parallel, to establish a baseline, the same analysis was performed on 20 randomly-selected genes from the same genomes from which each homolog originated, so that each genome serves as its own control for base composition, repeat content, and assembly quality. Moreover, to ensure annotation uniformity, genomes from the curated RefSeq database were analyzed separately from those found in GenBank.

We first interrogated the results with an initial count-based analysis of the distribution of repeat footprints (the number of nucleotides covered by repeat copies). As expected, a small fraction of randomly-selected genes had nearby repeats, ∼6% within 200 nucleotides of the gene (Figure 5D, top panel, dashed lines), likely owing to the presence of repetitive regulatory elements. However, most of these repeat footprints are short, and the fraction of repeat-bearing flanks surrounding these random genes declines steeply with increasing footprint size. In contrast, a greater fraction of *staN* homolog flanks harbor detectable repeats (21.3% of HTH+ flanks against 5.9% of background flanks, 3.6-fold; 6.5% of HTH++ flanks against 4.9%, 1.3-fold), and although the general trend of larger repeat footprints being rarer also holds true, the fraction of repeat-bearing flanks of *staN* homologs diminishes much less steeply with increasing footprint size. The result is that the ratio of these fractions (with repeat-bearing *staN* homolog flanks in the numerator) yields a fold enrichment value that increases with increasing footprint size (Figure 5D, bottom panel). This enrichment is most prominent in flanks bearing tandem direct repeats (TDR) surrounding HTH+ homologs, culminating in a peak enrichment of about 25-fold for flanks with at least 130 nucleotides of tandem repeat DNA.

To determine whether there is a statistically significant association between HTH class (HTH+ or HTH++) and the presence of repeats in the surrounding DNA relative to background genes from the same genomes, we built a rigorous Bayesian model using cmdstanr (an R package) with weakly informative regularizing priors. As an additional check, we also repeated the analysis and acquired point estimates using a maximum-likelihood fit of the same model, without priors (glmmTMB R package). The data showed that direct repeats (but not inverted repeats) are indeed enriched within 200 nucleotides (upstream and downstream) of HTH+ homologs compared to randomly-selected genes in the same genome: 7.9-fold for tandem and 7.4-fold for spaced direct repeats (Figure 5E). In addition, HTH+ homologs are more likely to lie proximal to direct repeats than HTH++ homologs. Our DPAM analysis also resolved the HTH domains into ECOD families, and a per-family breakdown showed a similar trend with few exceptions: the same HTH family is more likely to reside nearby repeats when it is a homolog’s only HTH and much less so in an HTH++ arrangement with other families (Figure 5F). We also investigated the spatial distribution of repeat enrichment and found that repeats are most likely to reside within 200 nucleotides of homologs, particularly in the upstream region of HTH+ homologs (12.4-fold for tandem and 10.6-fold for spaced direct repeats, Figure 5G).

We finally investigated the length distribution of repeating units and nucleotide composition of homolog loci. The data showed a periodicity to repeat length, with peaks at multiples of 10-12 nucleotides (Figure 5H), approximately one turn of the B-form DNA helix. Notably, nearly half of the HTH+ homologs with upstream repeats bear a tandem repeat array with a 10–12-nucleotide unit, 54 times the background rate, and 9.0% bear exactly 11-nucleotide units. Because a repeat unit length equal to an integer multiple of a single helical turn leaves successive units in the same rotational register, we can infer that proteins bound to adjacent units would be positioned on the same face of the helix. We also analyzed the nucleotide composition of StaN homolog flanks and coding regions, specifically the third codon positions, which are the least constrained by the protein, and thus show a gene’s compositional history most clearly^44,45^. This analysis revealed that coding regions are significantly more AT-rich than those of randomly-selected genes, with flanks also showing meaningful AT enrichment (Figure 5I). This compositional signature is a characteristic of mobile elements, such as phages, plasmids and insertion sequences^46^, suggesting that many StaN homologs may reside on mobile genetic elements.

### Distant StaN homologs exhibit toxicity that is mitigated by flanking DNA

We next sought to determine whether StaN homologs are functionally conserved. We selected a set of distant StaN homologs from diverse backgrounds with a variety of domain architectures and genetic contexts (Figure 6A). The selected homologs share 23-36 % identity across 65-89 % of their amino acid sequences in pairwise comparisons with StaN. To test for toxicity, each homolog was inserted downstream of the aTc-inducible promoter of pTET for expression in *S. aureus* and the arabinose-(ara-) inducible promoter of pBAD for expression in *E. coli*. Strains bearing the plasmids were grown in the presence of appropriate inducer, and colony counts were determined. The results showed that five of the homologs are highly toxic when expressed in one or both backgrounds (Figure 6 B and C). These homologs originate from diverse species, and notably, not all of them have flanking repeats.

**Figure 6.**
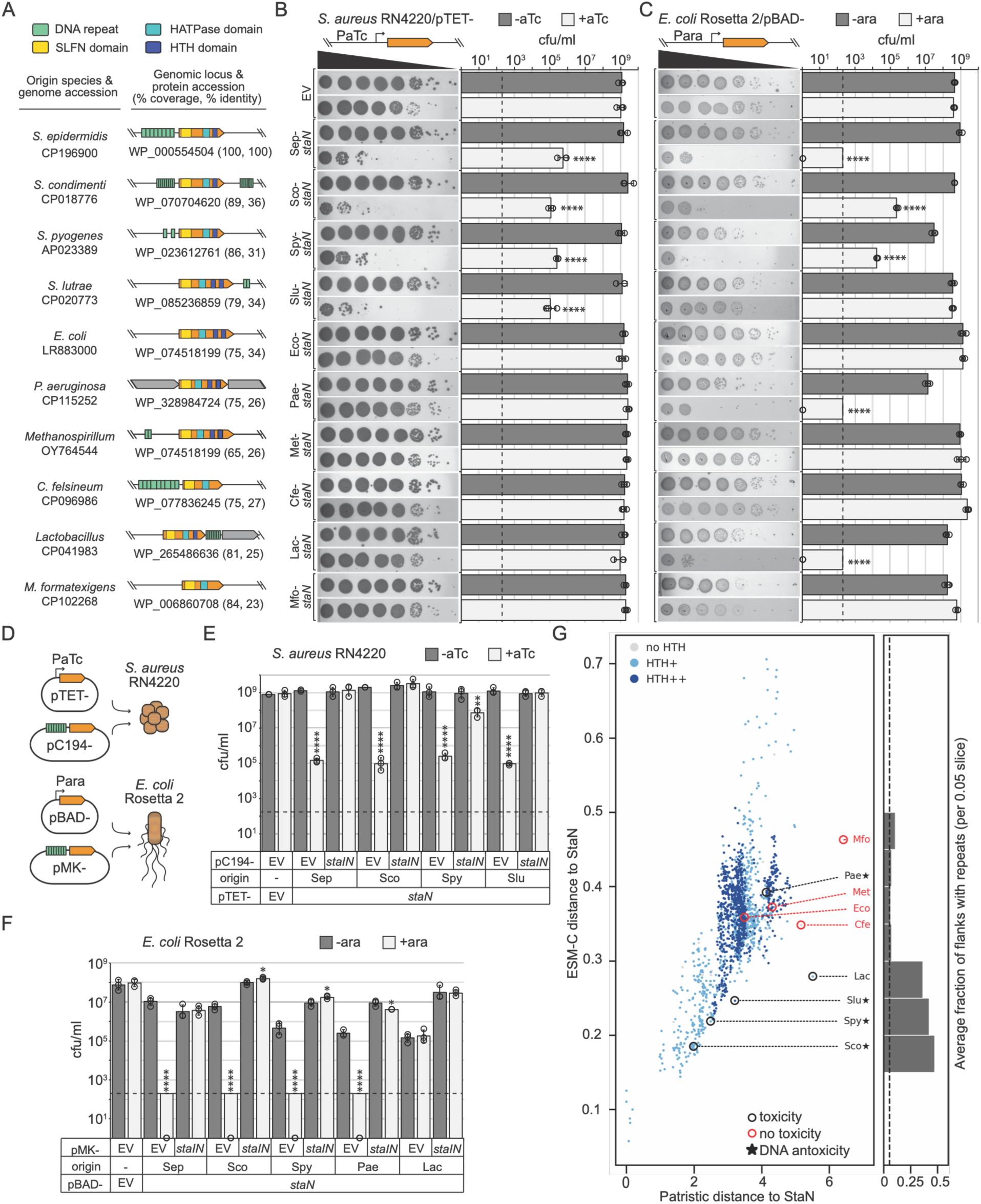
Diverse StaN homologs exhibit toxicity that is mitigated by flanking DNA. **(A)** StaN and nine selected homologs with respective domains and genomic contexts. **(B)** and **(C)** Cell toxicity assays with *staN* homologs in indicated strains. 10-fold dilutions of cells bearing indicated plasmids were plated +/-inducer (anhydrotetracycline (aTc) or arabinose (ara)). Average colony-forming units per milliliter (cfu/ml) from three technical replicates (+/- SD) and representative plate pictures are shown. Dotted line, limit of detection. Asterisks indicate p values < 0.0001(****) from two-tailed Student’s t-tests **(D)** Schematic of dual-plasmid cell toxicity assay. **(E)** and **(F)** Dual-plasmid toxicity assays with *staN* homologs. 10-fold dilutions of cells bearing indicated plasmids were plated +/- inducer. Average cfu/ml are shown from three technical replicates (+/- SD), and data is representative of two independent trials. Dotted line, limit of detection. Asterisks indicate p values < 0.05 (*), < 0.01(**), and < 0.0001(****) from two-tailed Student’s t-tests. **(G)** Distribution of StaN homolog cluster representatives plotted according to phylogenetic (patristic) and ESM-C embedding distances from StaN.

Next, we tested whether adding the coding region plus flanking DNA on a second plasmid *in trans* would mitigate toxicity (Figure 6D). As controls, strains harboring the empty vector (EV) versions of the second plasmid were included to rule out the possibility that the vector backbones themselves might interfere with toxicity. The results showed that the toxic phenotype persisted in all but one of the homologs (from *Lactobacillus*) in the presence of corresponding EVs, and notably, for the toxic homologs, the corresponding flanking DNA neutralized toxicity (Figure 6 E and F). These include the HTH++ *P. aeruginosa* homolog that shares 26% sequence identity with StaN and has no detectible flanking repeats. Altogether, these data demonstrate that distant StaIN systems from diverse bacteria maintain functional conservation, and flanking repeats may not be necessary to mitigate toxicity.

The results above suggest that amino acid sequence conservation is a weak predictor of function. In contrast, protein language models such as ESM-C (Evolutionary Scale Model-Cambrian) are trained precisely to relate structurally/functionally similar proteins in proximity to each other in their latent space. Thus, we compared the distances between StaN and its homologs inferred by ESM-C versus traditional phylogenetic analysis. We acquired embedding vectors for the 1,668 homolog cluster representatives (in Figure 5) using the 6 billion parameter ESM-C model, and after mean-pooling per residue vectors, calculated the cosine distance between all homologs and StaN. We also inferred pairwise distances between StaN and its homologs from the phylogenetic tree in Figure 5B. The results showed that while the homologs were spread across broad ranges of patristic and ESM-C distances from StaN, the representatives with demonstrated StaIN-like toxin/antitoxin function clustered more closely in ESM-C’s latent space (<0.3 distance to StaN) (Figure 6G). Moreover, the majority of homologs flanked by detectible repeats also fell below this threshold. These include the *Lactobacillus* system whose antitoxin activity could not be tested due to technical difficulties (Figure 6F). Together, these observations hint that the latter homolog may indeed exhibit DNA-based antitoxin activity.

## Discussion

Here we describe StaIN, a unique family of TA-like antiphage systems in which a toxic Slfn protein restrained by noncoding DNA provides population-level defense. Slfns are found in all domains of life and play broad roles in cellular growth and differentiation, cancer progression, and antiviral immunity^47^. In mammals, some Slfns restrict viral replication by cleaving tRNAs to limit viral protein synthesis^48–52^, and only recently has a similar function been ascribed to pSlfns, both as standalone antiphage defenses, and as effectors of Type III CRISPR immunity^28,29^. Nearly all pSlfns possess accessory domains that presumably act as phage sensors, with the majority (∼62%) comprising HATPase and/or HTH domains^28^. However, the previously-characterized examples bear sensor domains that are more rarely encountered—an immunoglobulin-like domain that recognizes phage tail assembly chaperones^28^, and a Rossmann-like domain that binds the Type III CRISPR-Cas second-messenger signaling molecule^29^. The mechanisms by which the predominant HATPase and HTH domains operate in antiphage defense remain unexplored.

Herein, we show that similarly to the first characterized pSlfns, StaN catalyzes metal-dependent tRNA cleavage via its SLFN domain; however, unlike the former, this activity requires ATP (Figure 3). Because the non-hydrolyzable analog ATPγS also supports cleavage, nucleotide occupancy rather than hydrolysis enables enzyme function (Figure 3 F and G). Moreover, StaN binds cooperatively to *staI* repeat DNA via its wHTH domain, and *staI* DNA (not RNA), is required to neutralize StaN’s toxicity (Figure 2). Thus, we propose a model in which StaN’s accessory domains exert two layers of control—the wHTH domain physically restrains StaN by tethering it to *staI* DNA, and the HATPase domain licenses tRNase activity by binding ATP (Figure 7).

**Figure 7.**
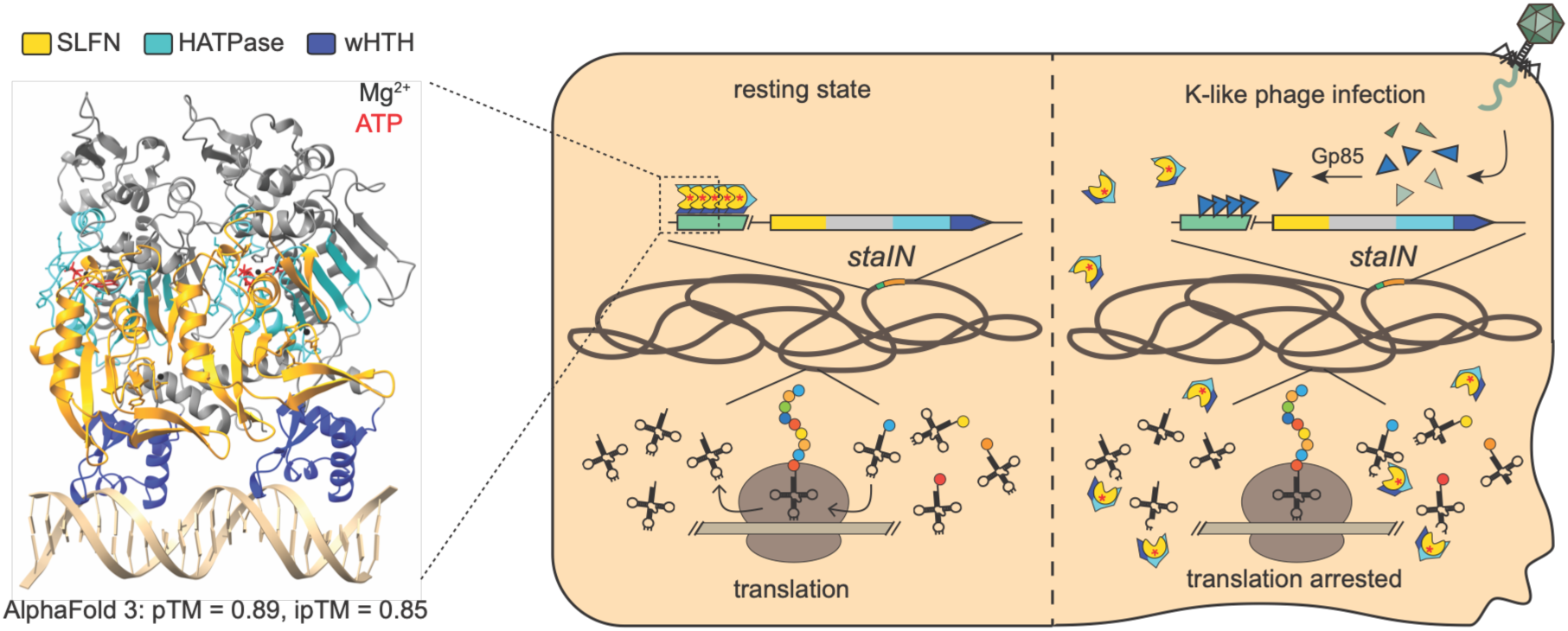
**Proposed mechanism of antiphage defense by StaIN and homologous systems**

The question then becomes—how does phage infection trigger StaN’s activation? ATP is among the most abundant metabolites in actively growing bacteria^53,54^, and since physiological ATP concentrations both support StaN’s toxicity in uninfected cells (Figure 2A) and activate tRNase activity in a purified system (Figure 3E), nucleotide occupancy is unlikely to be the infection-responsive step. Instead, we propose that release from *staI* DNA is the main driver of StaN’s activation. In agreement, extra copies of *staI* DNA provided *in trans* are sufficient to mitigate antiphage function in cells whose ATP availability remains otherwise unaltered (Figure 2D). Additionally, all phages that escape StaIN carry mutations in a hypothetical protein harboring a predicted DNA binding domain (ISP Gp85, Figure 4 A and B), and *in silico* analyses suggest that it too may bind to *staI* DNA (Figure S7D). Given that Gp85 expression promotes StaN’s toxicity (Figure 4C), we speculate that Gp85 displaces StaN from *staI* DNA upon phage infection. However, we cannot rule out other triggering mechanisms whereby Gp85 functions indirectly (see limitations below).

StaIN systems are widespread, present in diverse bacterial and archaeal phyla (Figure 5B), and we show that a set of distantly related representatives exhibit toxicity that is neutralized by flanking DNA (Figure 6). However, not all such flanks bear detectible DNA repeats, and more broadly, repetitive DNA is not a universally conserved feature of homologous StaIN systems (Figure 5 B and D). Considering the critical role of *staI* DNA in antitoxin activity, where neutralization requires the toxin to be bound by DNA, the dearth of homologs with detectible flanking repeats was initially surprising. However, our structure-based domain analysis, which recovered domains that sequence-based annotation had missed, revealed that a C-terminal HTH domain is retained in most homologs, and nearly half of all homologs carry multiple tandem HTH domains, typically belonging to different HTH families (Figure 5 C and F). Moreover, repeats are significantly more enriched near HTH+ homologs than their HTH++ counterparts (Figure 5 E and G). Among the tested representatives, the *Pseudomonas* homolog is particularly informative as it carries multiple HTH domains and has no detectable proximal repeats, yet its flanking DNA neutralizes toxicity (Figure 6F). We speculate that bearing multiple HTH domains from different families enables higher-affinity binding, and thus obviates the need for an array of DNA binding sites, provided that each HTH domain possesses a different binding specificity.

Finally, StaIN falls outside the current classification of TA systems, in which antitoxins are composed of either protein or RNA^4^. Additionally, our identification of Gp85 as a determinant of phage sensitivity to StaIN offers a concrete handle for designing phage-based therapeutics that are able to overcome immunity, while agents that mimic Gp85 and activate StaN would turn this resident defense effector against its host in an antibacterial strategy.

### Limitations of the study

The precise mechanism by which phage infection triggers StaIN immunity remains unclear. Our model is based on assays that measure StaN binding to *staI* DNA *in vitro* and the observation that *staI* DNA neutralizes StaN’s toxicity in cells. StaN occupancy of *staI* in cells has not been established, and we have not followed StaN localization during infection. The role of Gp85 is also inferred from *in silico* analyses, and it may act indirectly through host factor(s) rather than directly to trigger StaN’s deployment. Moreover, the exact function of Gp85 during phage infection and the basis for its toxicity in cells remains unknown^34^.

## Acknowledgements

A.H-A. would like to acknowledge funding for this project from the National Institutes of Health (NIH) Division of Intramural Research (1R01AI173022) and an Investigators in the Pathogenesis of Infectious Disease Award from the Burroughs Wellcome Fund (both awarded to A.H-A.). T.A.B. was funded by a Salyers Memorial Graduate Fellowship from the UIUC Microbiology Department.

## Author Contributions

Conceptualization (overall project), A.H-A.; Conceptualization (computational), B.A.; Data Curation, B.A.; Investigation, T.A.B., K.S., M.R.S., N.J.H., C.E.L., H.S.A.; Methodology, T.A.B., K.S., B.A.; Formal Analysis, T.A.B, B.A.; Software, B.A.; Visualization, T.A.B., B.A., and A.H-A.; Writing – Original Draft, A.H-A.; Writing – Review and Editing, T.A.B., K.S., B.A.; Supervision and Funding Acquisition, A.H-A.

## Declaration of Interests

The authors declare no competing interests.

## STAR Methods

### Key Resources Table

| **Reagent or Resource** | **Source** | **Identifier** |
|---|---|---|
| **Bacterial and virus strains** |  |  |
| *Staphylococcus epidermidis* LM1680 | Luciano Marraffini (Jiang et al.<sup>15</sup>) | Genbank: NC_002976.3 |
| *Staphylococcus aureus* RN4220 | Luciano Marraffini (Nair et al.<sup>14</sup>) | Genbank: NZ_CP150127.1 |
| *Staphylococcus aureus* USA300 | Tauqeer Alam (Diep et al.<sup>55</sup>) | Genbank: CP000255.1 |
| *Staphylococcus epidermidis* 6593473 | Stefania Stefani (This manuscript and Bongiorno et al.<sup>12</sup>) | Genbank: NZ_CP196900.1 |
| *Escherichia coli* DH5α | ATCC | Genbank: CP080399.1 |
| *Escherichia coli* Rosetta 2(DE3) | Novagen | Genbank: NZ_CP083274.1 |
| *Escherichia coli* Rosetta 2(DE3)pLysS | Novagen | Genbank: NZ_CP083274.1 |
| Bacteriophage ISP | Luciano Marraffini (Vandersteegen et al.<sup>56</sup>) | Genbank: NC_047720.1 |
| Bacteriophage Monmouth | This manuscript | Genbank: PZ600696 |
| Bacteriophage Lorac | Previous study (Marc et al.<sup>57</sup>) | Genbank: MH321492.1 |
| Bacteriophage Slyfox | This manuscript | Genbank: PZ600695 |
| Bacteriophage Pabna | Previous study (Culbertson et al.<sup>58</sup>) | Genbank: NC_048107.1 |
| Bacteriophage Merlin | This manuscript | Genbank: PZ600697 |
| Bacteriophage Twillingate | Previous study (Freeman et al.<sup>59</sup>) | Genbank: MH321491.1 |
| Bacteriophage Southeast | Previous study (Hill et al.<sup>60</sup>) | Genbank: OQ623150.1 |
| Bacteriophage Andhra | Previous study (Cater et al.<sup>61</sup>) | Genbank: KY442063 |
| Bacteriophage T4 | ATCC | Genbank: NC_000866.4 |
| **Chemicals, peptides, and recombinant proteins** |  |  |
| γ-[<sup>32</sup>P]-ATP | Revvity via Fisher Scientific | Part # BLU502H250UC |
| ATPγS | Enzo Life Sciences via VWR | Cat. # ALX-480-066 |
| *Escherichia coli* MRE 600 tRNA | Roche Diagnostics via Millipore Sigma | Cat. # 10109541001 |
| SUMO protease | MCLAB | Cat. # SP-100 |
| T4 polynucleotide kinase | New England Biolabs | Cat. # M0201L |
| DpnI | New England Biolabs | Cat. # R0176S |
| HisPur™ Ni-NTA Resin | Thermo Fisher | Cat. # M0201L |
| ACSVI7_RS00220 (StaN) | This manuscript | Genbank: NZ_CP196900.1_ACSVI7_RS00220 |
| QLX23_gp85 (ISP GP85) | This manuscript | Genbank: NC_047720.1_QLX23_gp85 |
| **Critical commercial assays** |  |  |
| EZNA Cycle Pure Kit | Omega Bio-tek via VWR | Cat. # 101318-892 |
| EZNA Plasmids DNA Mini Kit | Omega Bio-tek via VWR | Cat. # 101318-898 |
| Wizard® Genomic DNA Purification Kit | Promega Corporation via VWR | Cat. # A1120 |
| Wizard® DNA Cleanup Kit | Promega Corporation via VWR | Cat. # A7280 |
| PerfeCTa® SYBR® Green SuperMix | Quanta Biosciences via VWR | Cat. # 101414-150 |
| **Deposited data** |  |  |
| Illumina reads for ISP and escaper mutants | This manuscript | BioProject: PRJNA1522937 |
| **Oligonucleotides** |  |  |
| DNA oligonucleotides for PCR and Sanger sequencing | Eurofins MWG Operon | Table S6 |
| DNA oligonucleotides (PAGE purified) for biochemical assays | Eurofins MWG Operon | Table S8 |
| Synthetic constructs | Invitrogen GeneArt (via ThermoFisher Scientific) | Table S7 |
| **Recombinant DNA** |  |  |
| pCN57 | Luciano Marraffini (Charpentier et al.<sup>62</sup>) | Charpentier et al.<sup>62</sup> |
| P*crispr-cas* | Asma Hatoum-Aslan | Hatoum-Aslan et al.<sup>26</sup> |
| pC194 | ATCC (Horinouchi and Weisblum<sup>13</sup>) | Cat. # 37034 |
| pC194-based constructs (i.e. pC194-, multiple) | This manuscript | Table S6 |
| pKOR1 | Luciano Marraffini (Bae and Schneewind<sup>63</sup>) | Addgene Cat. # 133446 |
| pKOR1-based constructs (i.e. pKOR1-*staIN*) | This manuscript | Table S6 |
| pE194 | ATCC (Weisblum et al.<sup>64</sup>) | Genbank: M17811.1 |
| pE194-based constructs (i.e. pTarget, pTET-, multiple) | This manuscript and Samai et al.<sup>65</sup> | Table S6 |
| pBAD18 | Luciano Marraffini (Guzman et al.<sup>66</sup>) | Novopro Cat. # V001454 |
| pBAD18-based constructs (i.e. pBAD-, multiple) | This manuscript | Table S6 |
| pUC19 | NEB (Norrander et al.<sup>67</sup>) | Cat. # N3041S |
| pUC19-based constructs (i.e. pMK-, multiple) | Invitrogen GeneArt and this manuscript | Table S6 |
| pET28b | Novagen | Cat. # 69865 |
| pET28b derivatives (i.e. pET28b-, multiple) | This manuscript | Table S6 |
| **Software and algorithms** |  |  |
| BLASTp 2.17.0+ | Camacho et al.<sup>68</sup> | https://blast.ncbi.nlm.nih.gov |
| InterProScan (web) | Blum et al.<sup>27</sup> | https://www.ebi.ac.uk/interpro/search/sequence/ |
| InterProScan v. 5.77-108.0 (HPC) | Jones et al.<sup>69</sup> | https://github.com/ebi-pf-team/interproscan/releases/tag/5.77-108.0 |
| MMseqs2 (build 5d152c6) | Steinegger and Söding<sup>70</sup> | https://github.com/soedinglab/MMseqs2 |
| DPAM (v2.0) | Zhang et al.<sup>37</sup> | https://github.com/CongLabCode/DPAM |
| Tandem Repeats Finder 4.09 | Benson<sup>42</sup> | https://github.com/Benson-Genomics-Lab/TRF |
| Vmatch 2.3.1 | Fonseca et al.<sup>43</sup> | http://www.vmatch.de |
| Edlib 1.3.9 | Šošić and Šikić<sup>71</sup> | https://github.com/Martinsos/edlib |
| MAFFT 7.526 | Katoh and Standley<sup>39</sup> | https://mafft.cbrc.jp/alignment/software/ |
| trimAl 1.5.1 | Capella-Gutiérrez et al.<sup>40</sup> | https://github.com/inab/trimal |
| IQ-TREE 3.1.3 | Wong et al.<sup>41</sup> | https://iqtree.github.io |
| TreeViewer (v2.2.0) | Bianchini and Sánchez-Baracaldo<sup>72</sup> | https://treeviewer.org |
| CmdStan 2.39.0 and cmdstanr 0.9.0 | Carpenter et al.<sup>73</sup> | https://mc-stan.org |
| glmmTMB 1.1.14 | Brooks et al. | https://cran.r-project.org/package=glmmTMB |
| R 4.5.3 | R Core Team | https://www.r-project.org |
| Python 3.12 with NumPy, pandas and Biopython 1.87 | Python Software Foundation | https://www.python.org |
| UCSF ChimeraX 1.11.1 | Meng et al.<sup>74</sup> | https://www.cgl.ucsf.edu/chimera/ |
| ImageQuant TL v. 8.1 | GE Healthcare/Life Sciences | RRID: SCR_014246 |
| ImageJ 1.54g | Schneider et al.<sup>75</sup> | https://imagej.net/software/imagej/ |
| AlphaFold 3 (web) | Abramson et al.<sup>31</sup> | https://alphafoldserver.com/welcome |
| AlphaFold v. 3.0.1 (HPC) | Abramson et al.<sup>31</sup> | https://github.com/google-deepmind/alphafold3 |
| GraphPad Prism 11.0.0 | GraphPad Software, Inc. | https://www.graphpad.com |
| Mauve | Darling et al.<sup>76</sup> | https://darlinglab.org/mauve/mauve.html |
| Clinker | Gilchrist and Chooi<sup>77</sup> | https://cagecat.bioinformatics.nl/tools/clinker |
| bcl2fastq v. 1.8.4 | Illumina | https://support.illumina.com/sequencing/sequencing_software/bcl2fastqconversion-software/downloads.html |
| FastQC v. 0.11.9 | Babraham Bioinformatics | https://www.bioinformatics.babraham.ac.uk/projects/fastqc/ |
| SPAdes v. 3.15.3 | Bankevich et al.<sup>78</sup> | https://cab.spbu.ru/software/spades/ |
| Bandage v. 0.8.1 | Wick et al.<sup>79</sup> | https://rrwick.github.io/Bandage/ |
| bowtie2 v. 2.4.4 | Langmead and Salzberg | https://github.com/BenLangmead/bowtie2 |
| samtools v. 1.13 | Li et al.<sup>80</sup> | https://github.com/samtools/samtools/releases/ |
| igvtools v. 2.11.1 | Robinson et al.<sup>81</sup> | https://software.broadinstitute.org/software/igv/2.11.x |

### Resource Availability

#### Lead contact

Requests for further information and/or reagents and resources should be directed to and will be fulfilled by the lead contact, Asma Hatoum-Aslan.

#### Materials availability

Phages, strains, mutant derivatives, and constructs generated in this study will be made available upon written request to the lead contact.

#### Bacterial strains and growth conditions

*S. epidermidis* LM1680^15^, which is a mutant variant of *S. epidermidis* RP62a^82^ and *S. aureus* RN4220^14^ were generous gifts from Luciano Marraffini.

*S. epidermidis* 6593473 (this manuscript) was a generous gift from Stefania Stefani. *S. aureus* USA300^55^ was a generous gift from Tauqeer Alam. *S. epidermidis* strains were grown in Brain Heart Infusion (BHI, BD Diagnostics), *S. aureus* strains were grown in Tryptic Soy Broth (TSB, BD Diagnostics), *E. coli* strains were grown in Luria Bertani (LB) Broth (VWR), and *E. coli* Rosetta 2(DE3)pLysS strains were grown in Terrific Broth (TB, VWR) for protein purification. Bacterial growth media was supplemented with the following antibiotics when necessary: 10 μg/ml chloramphenicol (selection of pC194-based plasmids), 10 μg/ml erythromycin (selection of pTET-based plasmids), 15 μg/ml neomycin (selection of *S. epidermidis* strains), 30 μg/ml chloramphenicol (selection of *E. coli* Rosetta 2(DE3)pLysS plasmids), 50 μg/ml kanamycin (selection for pET28b-10HisSmt3-based and pMK-based plasmids), and 100 μg/ml ampicillin (selection of pBAD-based plasmids). All bacterial strains were grown at 37°C. Liquid cultures were grown with agitation at 180 rpm in an orbital shaker. Strains were regularly authenticated using PCR amplification and Sanger sequencing of unique regions or Oxford Nanopore whole-plasmid and amplicon sequencing (performed by Eurofins MWG Operon or Plasmidsaurus).

#### Bacteriophage propagation and enumeration

*S. aureus* phages (ISP, Monmouth, Lorac, Slyfox, and Pabna), and *S. epidermidis* phages (Merlin, Twillingate, Southeast, and Andhra), and *E. coli* phage T4 were grown on their host strains, *S. aureus* RN4220 and *S. epidermidis* LM1680, and *E. coli* DH5α. High-titer phage stocks were created by picking one thrice-purified phage plaque from a top-agar bacterial overlay and resuspending with 500 μl of TSB. Suspensions were pelleted for 2 minutes at 15,000 x *g*, and 200 μl of the resulting supernatant was added to 300 μl of host overnight culture, 5 mM CaCl2, and 7 ml of molten (55°C) Heart Infusion Agar (HIA, Hardy Diagnostics, 0.3 x concentration). Mixtures were then poured onto a Tryptic Soy Agar (TSA, BD Biosciences) plate and allowed to solidify on the benchtop for 10 minutes. After overnight incubation at 37°C, the phage/host top agar layer was scraped off the plate, combined with 20 ml of TSB, and vortexed for 5 minutes to release phages from the agar. The suspension was subjected to centrifugation at 10,000 x g for 10 minutes, and the resulting supernatant was passed through a 0.45 μm bottle filter to separate phage lysate from cell debris. Phage concentrations (titers) were calculated using a double-agar overlay method as previously described^61^. In brief, molten HIA (0.5 x concentration) was combined with 100 μl of overnight host culture and 5 mM CaCl_2_ and poured atop a TSA plate supplemented with 5 mM CaCl_2_. While the top agar overlay solidified for 10 minutes at room temperature, concentrated phage lysate was serially diluted in TSB (10^0^-10^-7^). 5 μl spots of serially diluted lysates were spotted on top of double-agar overlay plates and were allowed to dry before incubating at 37°C overnight. The phage titers in plaque-forming units per milliliter (pfu/ml) were calculated by dividing the number of plaques counted in most diluted spot by the dilution factor and multiplying by 200. Phage stocks were maintained at 4°C for short-term use and frozen with DMSO (10% final concentration) at-80°C for long-term storage. Phage stocks were regularly authenticated through PCR amplification and Sanger sequencing of unique phage genomic regions.

## Method Details

### Plate-based infection assays

All plate-based infection assays were performed by spotting phage serial dilutions atop a double-agar overlay plate and calculating phage titers, as described in the section above, at 37°C (unless indicated otherwise in the figure). Graphs show an average of three technical replicates (+/- standard deviation (SD)) from a representative of three independent trials.

### Constructing pC194, pKOR1, pE194-based plasmids

Gibson Assembly^83^ and inverse PCR were used to construct pC194-, pKOR1-, and pE194-based plasmids with the primers listed in Table S6. pC194^13^ was used as the backbone for all plasmids listed pC194-except for pC194-P*crispr-gfp*, which used p*crispr-cas*^26^, a derivative of pC194 that encodes the entire CRISPR-Cas10 system from *S. epidermidis* RP62a^82^. pKOR1^63^ was used as the backbone for pKOR1-*staIN*. pTarget^65^, a derivative of pE194^64^, was used as the backbone for all plasmids listed pTET-in this study. Both pC194 and pE194 are episomal plasmids with a medium-high copy number (10-20 copies/cell) in staphylococci^13,64^. pKOR1 was utilized to introduce a single copy of *staIN* into the staphylococcal genome through allelic replacement, as described below in the section titled, “Constructing single-copy knock-in strains.” All plasmids were introduced into *S. aureus* RN4220 through electroporation, as described in the next section, and plasmids were confirmed through PCR amplification and Sanger sequencing of insert regions (primers listed in Table S6) or Oxford Nanopore whole-plasmid/amplicon sequencing (performed by Eurofins MWG Operon or Plasmidsaurus). At least two independent transformants were confirmed by sequencing, and one of each construct was purified using an EZNA Plasmid Mini Kit (Omega Bio-tek) and introduced into *S. aureus* RN4220::*staIN*, S*. epidermidis* LM1680, or strains with existing plasmids through electroporation where indicated.

### Electroporation into staphylococci

Electrocompetent cells were prepared as described previously^84^. In brief, 10 ml of overnight culture was diluted to an optical density at 600 nm (OD600) = 0.5 with TSB and grown with agitation for 30 minutes at 37°C. Cells were cold-shocked for 10 minutes on ice with all further steps on ice or at 4°C. Following incubation, cells were pelleted at 4,000 x *g* for 10 minutes at 4°C and washed twice with an equal volume of ice-cold water. Cells were then washed with 10% glycerol at 1/20 and 1/25 of the volume of the original culture. Finally, cells were resuspended in 1/200 of the volume of the original culture, divided into 50 μl aliquots, and stored at-80°C. To electroporate competent cells, purified plasmids, Gibson assembled constructs, or ligated inverse PCR constructs were first dialyzed on 0.022 μm filters for 20 minutes. During the dialysis step, competent cells were thawed on ice and the benchtop for 5 minutes each before pelleting at 5,000 x *g* for one minute. The resulting pellet was resuspended in 50 μl of 10% glycerol supplemented with 500 mM sucrose before adding the entire volume of dialyzed plasmid or construct. The resulting suspension was moved to a 2 mm electroporation cuvette (VWR) and pulsed at 29 kV/cm, 100W, and 25 mF with a GenePulser Xcell instrument (Bio-Rad). Cells were recovered in 1 ml of TSB supplemented with 500 mM sucrose and incubated with agitation at 37°C for two hours. 200 μl of the recovered cells was plated on TSA with the appropriate antibiotics and incubated overnight at 37°C. Colonies resistant to the selected antibiotics were recovered and confirmed for plasmid presence the following day.

### Constructing single-copy knock-in strains

Allelic exchange utilizing pKOR1^63^ was performed as described previously^85^ with minor modifications. In brief, allelic exchange was used to introduce *staIN* into the chromosome of *S. aureus* RN4220. RN4220 genomic DNA was used as the template for PCR amplifying 1 kb upstream and downstream homology arms, and *S. epidermidis* 6593473 genomic DNA was used as template for amplifying *staIN.* pKOR1 was used as template for amplifying the pKOR1 backbone. Single overlap extension PCR^86^ was used for combining *staIN* and upstream/downstream fragments together for two-piece Gibson assembly^83^ with the pKOR1 backbone (all primers listed in Table S6). Assembled products were electroporated into *S. aureus* RN4220 and recovered cultures were plated on TSA supplemented with chloramphenicol and incubated at 30°C for 48 hours. Colonies were screened for the presence of the plasmid via PCR and confirmed clones were selected for subsequent steps. Clones were struck out on pre-warmed TSA plates with chloramphenicol and grown overnight at 43°C. This step was repeated a second time, and junction PCR confirmed the presence of the integrated plasmid. Confirmed colonies were passaged at 30°C in TSB for 3 days to facilitate excision and loss of the plasmid. 1:10,000 dilutions were made of the final passage and plated on TSA with 50 ng/μL anhydrotetracycline. Colonies were replica-plated on TSA supplemented with chloramphenicol and TSA to detect if the plasmid had been lost. Clones that could not grow on chloramphenicol were then screened using primers outside of the homology arms to check for the presence of the insert. Clones with a larger PCR product than the ancestral *S. aureus* RN4220 were then streak-purified and sequenced to confirm the insertion of *staIN*. Five independent RN4220::*staIN* knock-in strains were confirmed with PCR and one strain was confirmed with PCR and Sanger sequencing.

### One-step phage growth curves

One-step phage growth curves were performed as described previously^61^ with modifications. In brief, 20 ml of *S. aureus* RN4220, RN4220::*staIN*, and RN4220 bearing pC194-*staIN* or empty vector were grown to an OD600 = 0.5, supplemented with 1 mM CaCl_2_, and infected with a volume of phage lysate to reach a multiplicity of infection (MOI) of 0.1. Phages were allowed to adsorb to the host cells for 10 minutes at room temperature before pelleting at 4,000 x *g* for 5 minutes via centrifugation. Cells were washed with 20 ml of TSB before final resuspension in 20 ml of TSB. 1-ml aliquots of resuspended cells were incubated at 37°C, removed at 5-minute intervals, and pelleted via centrifugation. Supernatants were passed through 0.45 μm syringe filters to separate phage from bacterial cells, and dilutions were spotted atop double-agar overlays, as described in the section titled, “Bacteriophage propagation and enumeration.” Graphs show an average of three technical replicates (+/-SD) from a representative of three independent trials.

### Adsorption assays

Phage adsorption assays were performed as described previously^17^ with modifications. In brief, 10 ml of *S. aureus* RN4220 and RN4220::*staIN* were grown to an OD600 = 0.5, supplemented with 1 mM CaCl_2_, and infected with phage to reach an MOI of 0.1. Phages were allowed to adsorb to host cells for 10 minutes at room temperature before the cells were pelleted via centrifugation for 5 minutes at 4,000 x *g*. Supernatants containing unadsorbed phages were passed through 0.45 μm syringe filters, and dilutions were spotted atop double-agar overlays, as described in the section titled, “Bacteriophage propagation and enumeration.” A control sample with phage and no bacterial cells was enumerated and designated as the total number of phages added. To calculate the number of phages adsorbed, the number of unadsorbed phages was subtracted from the total number of phages added. Graphs show an average of three technical replicates (+/-SD) from a representative of two independent trials.

### Phage infection time course and quantitative PCR

Phage infection time course experiments were performed as described previously^85^ with modifications. In brief, 80 ml of *S. aureus* RN4220 bearing pC194-*staIN* or the empty vector were grown to an OD600 = 0.5, supplemented with 5 mM of CaCl_2_, and infected with phage to reach an MOI of 0.5. Phages were allowed to adsorb for 10 minutes at room temperature. Host/phage mixtures were divided into four 20-ml aliquots and incubated with agitation at 37°C for 0, 10, 20, or 30 minutes. Cells were then pelleted via centrifugation for 5 minutes at 4,000 x *g* and washed twice with 1X phosphate-buffered saline (PBS). The resulting pellets were resuspended in 200 μl of sterile water supplemented with 20 μg of lysostaphin (AmbiProducts) and 5 mM MgCl_2_ and incubated at 37°C for two hours. Total genomic DNA was extracted using the Wizard® Genomic DNA Purification Kit (Promega Corporation). To prepare 25-μl qPCR reactions, 500 ng of total DNA extracted from each time point, 0.4 nM of phage or host-specific primers (Table S6), and 1X PerfeCTa SYBR Green SuperMix (Quanta Biosciences) were combined. Standard reactions composed of 10^2^–10^9^ DNA molecules were prepared with phage ISP or *S. aureus* RN4220 extracted DNA. DNA templates were amplified using a CFX Connect Real-Time PCR Detection System (Bio-Rad) using the following protocol: one cycle, 95°C for 3 minutes; 40 cycles, 95°C for 10 seconds and 55°C for 30 seconds. Phage DNA values were normalized against host values, and the 0-minute time point was set to one to compare to the DNA abundance increase in later time points. Relative DNA abundance was determined using the following equation from the “Real-Time PCR Handbook” (ThermoFisher Scientific): Fold Difference = (E_target_)^Ct_target^ / (E_normalizer_)^ΔCt_normalizer^, where E = 10^(-1/slope)^, Ct_target = Ct_target_calibrator_-Ct_target_samples_, and ΔCt_normalizer = Ct_normalizer_calibrator_-Ct_normalizer_samples_. Graphs show an average of three technical replicates (+/-SD) from a representative of two independent trials.

### Cell growth dynamics assays

Cell growth assays were performed as described previously^17^. In brief, *S. aureus* RN4220, RN4220::*staIN*, and RN4220 bearing pC194-*staIN* or empty vector were diluted 1:100 in TSB, supplemented with 5 mM CaCl_2_, and infected with phage to reach MOI values of 0, 0.05, 1, and 5. 200 μl of host/phage mixtures were distributed in triplicate into a 96-well microtiter plate and incubated with agitation at 37°C (unless indicated otherwise in the figure) in an Infinite 200 Pro (Tecan). OD600 measurements were taken every 15 minutes for a period of 600 minutes. Graphs show an average of three technical replicates (+/-SD) from a representative of three independent trials.

### Cell toxicity assays

For cell toxicity assays, overnight cultures of *S. aureus* and *E. coli* bearing pTET or pBAD-based plasmids were serially diluted in TSB (10^0^ - 10^-7^). 5 μl spots of serially diluted cells were spotted on top of TSA plates supplemented with either 30 ng/ml anhydrotetracycline or 10 mg/ml arabinose and allowed to air dry before incubating at 37°C overnight. The cell titers in colony-forming units per milliliter (cfu/ml) were calculated by dividing the number of colonies counted in most diluted spot by the dilution factor and multiplying by 200. Graphs show an average of three technical replicates (+/- SD) from a representative of three independent trials.

### GFP reporter assay

For GFP reporter assays, overnight cultures of *S. aureus* RN4220 bearing pC194-*gfp*-based plasmids were diluted 1:100 in TSB. 200 μl of culture was distributed in triplicate into a 96-well microtiter plate and incubated with agitation at 37°C in an Infinite 200 Pro (Tecan). OD600 and fluorescence measurements (excitation/emission wavelengths: 485/535 nm, gain: 114) were taken every 15 minutes for a period of 600 minutes. Relative Fluorescence Units (RFUs) were normalized to cells bearing the empty vector. Graphs show an average of three technical replicates (+/-SD) from a representative of three independent trials.

### Construction of pBAD, pET28b-10His-Smt3, and pUC19-based plasmids

Gibson Assembly^83^ and inverse PCR were used to construct pBAD-, pET28b-10His-Smt3-, and pUC19-based plasmids with the primers listed in Table S6. pBAD18^66^ was used as the backbone for all plasmids listed pBAD-. pET28b (Novagen) was used as the backbone for plasmids listed pET28b-10His-Smt3-. pMK-RQ (Invitrogen GeneArt synthetic constructs), a derivative of pUC19^67^ with kanamycin resistance, was used as the backbone for plasmids listed pMK-. pBAD18 and pET28b are episomal plasmids with a low-medium copy number^66,87,88^, while pUC19 maintains a high copy number in *E. coli*^88,89^. Backbones were amplified using PCR, and supercoiled template plasmids were degraded with DpnI (NEB) before Gibson assembly or inverse PCR, phosphorylation, and ligation. All plasmids were introduced into chemically-competent *E. coli* DH5α cells, as described in the next section, and plasmids were confirmed through PCR amplification and Sanger sequencing of insert regions (primers listed in Table S6) or Oxford Nanopore whole-plasmid/amplicon sequencing (performed by Eurofins MWG Operon or Plasmidsaurus). At least two independent transformants were confirmed by sequencing, and one of each plasmid was purified using an EZNA Plasmid Mini Kit (Omega Bio-tek) and introduced into *E. coli* Rosetta 2(DE3), Rosetta 2(DE3)pLysS, or strains with existing plasmids where indicated.

### Transformation of E. coli

*E. coli* DH5α, Rosetta 2(DE3), and Rosetta 2(DE3)pLysS chemically-competent cells were prepared as described previously^17^. Overnight cultures were diluted 1:100 in 50 ml of LB and grown at 37°C with agitation to an OD600 = 0.5. Cell growth was stopped by placing cultures on ice for 10 minutes, and all future steps were performed on ice or at 4°C. Cells were pelleted via centrifugation for 10 minutes at 4,000 x g and the resulting pellet was resuspended in 5 ml of Transformation and Storage Solution (TSS) buffer (85% LB medium, 10% (w/v) polyethylene glycol (PEG) 8000, 5% (v/v) DMSO, and 50 mM magnesium chloride). Cells were divided into 50-μl aliquots and stored at-80°C. For transformation of *E. coli* cells, chemically-competent cells were thawed on ice for 10 minutes. 5 μl of Gibson assembly or inverse PCR product and 1 μl of purified plasmid was mixed with the cells and allowed to incubate on ice for 30 minutes. Mixtures were then heat-shocked at 42°C for 30 seconds and immediately placed back on ice for 2 minutes. Cells were recovered by adding 1 ml of LB and incubating at 37°C with agitation for one hour. 100 μl of transformed cells was spread on an LB-agar plate with the appropriate antibiotics and incubated overnight at 37°C. Colonies resistant to the selected antibiotics were recovered and tested for plasmid presence the following day.

### Purification of recombinant StaN

Recombinant StaN and mutant variants were overexpressed from *E. coli* Rosetta 2(DE3)pLysS cells bearing pET28b-10His-Smt3-based plasmids and purified as described previously^85^ with modifications. Briefly, one liter of TB supplemented with the appropriate antibiotics was inoculated with 10 ml of overnight culture. Cells were grown at 37°C with agitation until the OD600 reached 0.5-0.6, and protein expression was induced with 0.3 mM isopropyl-1-thio-b-d-galactopyranoside (IPTG) and 2% ethanol. Cells were allowed to grow for an additional 14-16 hours at 17°C with agitation. After pelleting via centrifugation, harvested cells were washed with ice-cold PBS and resuspended in 30 ml of Buffer A (50 mM Tris-HCl pH 8, 1.25 M NaCl, 200 mM LiSO_4_, 10% sucrose, and 15 mM imidazole) supplemented with one EDTA-free protease inhibitor tablet (Roche), 0.1% Triton X-100, and 0.1 mg/ml lysozyme, and all further steps were carried out at 4°C or on ice. Cells were rotated for one hour before the mixture was sonicated. Lysed cell debris was separated from soluble materials via centrifugation and filtration, and filtrate was incubated with 4 ml of HisPur™ Ni^2+^-NTA-agarose resin (ThermoFisher) that was pre-equilibrated with Buffer A for one hour. Following incubation, the resin was pelleted via centrifugation, washed with Buffer A, and loaded onto a 5-ml gravity column (G-Biosciences). The loaded column was further washed with 25 ml of Buffer A before eluting proteins in a stepwise fashion. Stepwise elution solutions contained increasing concentrations of imidazole (50, 100, 200, and 500 mM) in IMAC buffer (50mM Tris-HCl pH 8, 250 mM NaCl, 10% glycerol), and 3 ml of each concentration was used for elution. The proteins eluted from the 200 mM and 500 mM solutions were then pooled and mixed with 1000 U of SUMO protease and salt-free buffer (MCLAB) to remove the 10His-Smt3 tag, loaded into a 10K MWCO Slide-A-Lyzer® dialysis cassette (ThermoFisher), and dialyzed for 3 hours. Following tag clipping, the dialyzed sample was mixed with 1 ml of pre-equilibrated of HisPur™ Ni^2+^-NTA-agarose resin and rotated for one hour before it was loaded onto a 5-ml gravity column. Tag-free protein was collected in the flow-through and pooled for concentration using a 10K MWCO centrifugal filter (PALL). Protein samples were further purified by size exclusion chromatography using a Superdex® 200 increase 1/300 GL column (Cytiva). Resulting fractions were resolved on a 10% SDS-PAGE gel, stained with Coomassie G-250, and aliquots with the expected size and highest peak were pooled and concentrated for downstream tRNA cleavage and DNA-binding assays. Pooled protein samples were also resolved on a 10% SDS-PAGE gel and stained with Coomassie G-250. Concentrations were determined using a NanoDrop™ spectrophotometer (ThermoFisher). Three independent purifications of the wild-type (WT) StaN and two independent purifications of the mutant variants were conducted.

### tRNA cleavage assays

tRNA cleavage assays were performed with 1 μM of *E. coli* MRE 600 tRNA (Roche) and 0.5 μM of WT StaN or mutant variants in 10 μl reactions with nuclease buffer (25 mM Tris-HCl, pH 8 and 2 mM DTT) supplemented with 2 mM MgCl_2_ (or MnCl_2_, CaCl_2_, EDTA where indicated) and 1 mM ATP (or ATPγS/H_2_O where indicated). Experiments were incubated at 37°C for 40 minutes or 0, 10, 20, 40, and 80 minutes. Reactions were stopped by incubating on ice for 5 minutes, followed by treatment with 10 μg of proteinase K at room temperature for 20 minutes. Samples were then mixed with an equal volume of 95% formamide loading buffer and resolved on a 12% Urea-PAGE gel stained with SYBR™ Gold (ThermoFisher).

Quantification of tRNA was performed using densitometric analysis from ImageJ version 1.54g^75^.To determine percentage of tRNA cleaved, the density of the uncut tRNA substrate from the experimental samples was divided by that of the negative control (no protein added). Representative images of three independent trials are displayed, and the tRNA cleavage graph displays an average of three independent trials (+/-SD).

### DNA binding assays

For DNA binding assays with 75-nt dsDNA, single-stranded DNA (ssDNA) substrate (Table S8) was labeled on its 5’-end by incubating with γ-[^32^P]-ATP (Revvity) and T4 polynucleotide kinase (NEB) for 30 minutes at 37°C. The ssDNA was purified using a G25 column (IBI Scientific) and mixed at a 1:2 molar ratio with unlabeled complementary ssDNA. The mixture was heated at 95°C for 5 minutes, followed by incubation at room temperature for at least 3 hours. 50 nM of the double-stranded DNA (dsDNA) probe was then mixed with increasing concentrations of WT StaN (40, 200, 400, 800, 1200, 1600 2000, 4000 nM) and incubated for 15 minutes at room temperature before resolving on a 10% Native-PAGE gel. The gel was dried, exposed to a storage phosphor screen, and visualized using an Amersham™ Typhoon™ biomolecular imager. To quantify bound and free dsDNA, Amersham™ ImageQuant™ TL v.8.1 software (GE Healthcare/Life Sciences) was used for densitometric analysis (minimum slope = 50 and edge parameter = fixed at 16). Percentage of DNA bound was calculated using the following formula: density of substrates shifted to bound state divided by the sum of densities in the bound and free states. To calculate the dissociation constants (Kd), percent DNA bound was plotted versus protein concentration, and a specific binding with Hill slope equation was fit to these data using GraphPad Prism 11.0.0. For DNA-binding assays with 500-nt dsDNA, substrates were amplified via PCR (primers listed in Table S6) and purified using the EZNA Cycle Pure Kit (Omega Bio-tek). 0.3 μM of dsDNA substrate was then mixed with 0.6 μM of either WT StaN or ΔWH (winged-helix) mutant variant and incubated for 15 minutes at room temperature before resolving on a 2% agarose gel. The gel was then stained with SYBR™ Gold (ThermoFisher). For the 75-nt dsDNA-binding experiment, a representative image and a graph of two independent trials is shown. For the 500-nt dsDNA experiment, a representative image of three independent trials is shown.

### Phage infection time course and DNA cleavage assays

Phage infection time course experiments were performed as described previously^85^ with modifications. Briefly, 40 ml of *E. coli* DH5α and *S. aureus* RN4220 were grown to an OD600 = 0.5, supplemented with 5 mM of CaCl_2_, and infected with phage T4 or ISP to reach an MOI of 10. Phages were allowed to adsorb for 10 minutes at room temperature. Cells were divided into four 10-ml aliquots and incubated with agitation at 37°C for 0, 5, 10, or 15 minutes. Cells were then pelleted via centrifugation for 5 minutes at 4,000 x *g* and washed twice with 1X PBS. The resulting *S. aureus* pellets were resuspended in 100 μl of sterile water supplemented with 20 μg of lysostaphin (AmbiProducts) and 5 mM MgCl_2_ and incubated at 37°C for two hours. The resulting *E. coli* pellets proceeded directly to DNA extraction. Total genomic DNA was extracted using the Wizard® Genomic DNA Purification Kit (Promega Corporation). Following extraction, 250 ng of DNA was resolved on a 0.6% agarose gel and stained with SYBR™ Gold (ThermoFisher).

### Isolation and amplification of ISP escaper phages

ISP escaper phages were isolated by spotting 10-fold serial dilutions (10^0^ - 10^-7^) of phage ISP atop a double-agar overlay plate with *S. aureus* RN4220 bearing pC194-*staIN.* Plates were incubated overnight at 37°C, and 12 individual plaques were picked with a sterile pipette tip from two independent trials. Plaques were resuspended in 500 μl of TSB and spotted atop a double-agar overlay plate with *S. aureus* RN4220 bearing pC194-*staIN*. This process was repeated thrice. Following purification, large-scale propagation was performed as described in the section titled, “Bacteriophage propagation and enumeration.”

### Genomic DNA extraction of ISP escaper phages

Genomic DNA was extracted from high-titer (at least 1 x 10^8^ pfu/ml) phages as described previously^61^ with minor modifications. 40 ml of phage lysate was split between two tubes and digested with 10 μg/ml of DNase I and RNase A for 30 minutes at 37°C. Lysates were then mixed with 10 ml of precipitant solution (30% (wt/vol) PEG 8000 and 3 M NaCl) and incubated at 4°C overnight. Precipitated phage solutions were pelleted via centrifugation for 10 minutes at 10,000 x *g* and 4°C. Pellets were combined and resuspended in 250 μl of 5 mM MgSO_4_ and further digested with 400 μg/ml of proteinase K and 10 mM EDTA (pH 8.0) at 50°C for 30 minutes. The phage suspensions were allowed to cool to room temperature, mixed with resin from the Wizard® DNA Cleanup Kit (Promega), and inverted 5-6 times. The resins were then loaded onto the provided minicolumns, washed with 2 ml of 80% isopropanol, and dried via centrifugation at 13,000 x *g* for 2 minutes. Finally, phage DNA was eluted by adding 100 μl of distilled H_2_O at 80°C to the minicolumns, and concentrations were measured using a NanoDrop™ spectrophotometer.

### DNA sequencing and analysis of ISP escaper phages

ISP escaper phage DNA library preparation and sequencing was performed as described previously^17^ by SeqCenter with the Illumina DNA Prep kit and Illumina NextSeq 2000. Adaptor and index sequence removal was performed by bcl2fastq v. 1.8.4 (Illumina), and data quality was checked with FastQC v. 0.11.9 (Babraham Bioinformatics). To assemble reads from ancestral ISP, SPAdes v. 3.15.3^78^ was used in isolate mode with the kmer values of 21, 33, 55, 77, and 99. Bandage v. 0.8.1^79^ was then used to visualize the uploaded genome assembly graph and extract the high coverage phage genome as a FASTA file. After the ancestral ISP FASTA file was indexed using bowtie2 v. 2.4.4^90^, escaper phage sequencing reads were recruited to the indexed sequence with the default alignment and scoring parameters. The resulting Sequence Alignment/Map (SAM format) file was converted to a binary (BAM) format and sorted with samtools v. 1.13^80^. The indexed file was then used for coverage depth analysis with igvtools v. 2.11.1^81^. The resulting file was further analyzed to calculate the fraction of escaper phage reads that match to ancestral ISP at every nucleotide and the corresponding depth in coverage, which was plotted for all twelve unique escaper phages.

### Homolog search

We searched the RefSeq protein database with BLASTp 2.17.0+, using full-length StaN (WP_000554504.1, 496 amino acids) as the query (BLOSUM62, gap costs 11/1, low-complexity filter off, default E-value cutoff 0.05). For our purposes, in order to cast a wide net, we chose BLASTp search over iterative profile methods such as PSI-BLAST or JackHMMER to avoid over constraining the search. The search returned its maximum of 5,000 hits before reaching the E-value cutoff (weakest hit E = 1.6 × 10⁻¹⁸), which suggests that the StaN family likely extends beyond the sample set analyzed here. We set aside three hits that are 60-amino-acid fragments with no detectable domain. The nine homologs tested experimentally alongside StaN had been chosen from a phylogeny of an earlier, smaller BLAST search; five were already among the new hits, and we manually added the remaining four that fell beyond the 5,000-hit limit, resulting in the 5,001 homologs. We used NCBI Identical Protein Groups (IPG) to link each one to the genome records that encode it.

### Clusters and domain architectures

Many hits were near-identical copies of one protein from closely related strains, so we clustered the homologs with MMseqs2^70^ at 95% identity and 95% coverage, resulting in 2,287 clusters. We predicted the structures of the cluster representatives and (separately) the ten tested proteins (including StaN) with AlphaFold 3^31^ and used DPAM^37^ to split each structure into ECOD^38^ domains, resulting in 2293 solved structures; cluster members without a prediction of their own took the domain architecture of their representative (overall, cluster members were median 98.4% identical). Every domain in ECOD’s HTH group were counted as an HTH domain, named by its ECOD family, if available, and homologs were grouped by their number of HTH domains: none, one (HTH+), or two or more (HTH++). Based on these architectures, we retained only those homologs that start with a methionine, carry complete Schlafen and RecG (HATPase) domains, and are not an exact fragment of a longer homolog that carries a domain they lack. These “intact” homologs of each cluster were analyzed together, leaving 1,668 clusters with 4,036 homologs. To compare structure-based with sequence-based domain calls, we ran InterProScan 5.77-108.0^69^ (default member databases) on all 5,001 homologs, merged overlapping signatures of one domain into a single locus, and compared each representative’s InterProScan HTH count with its DPAM count. To see whether HTH domains remain a common feature even among the more divergent homologs, we split the intact homologs into quarters by BLAST bit score and counted the share of each quarter carrying an HTH domain. Even though a single HTH domain makes up only about 15% of a homolog’s length, we observed that even among the quarter of homologs least similar to StaN (median 30% identity to StaN), 96% carry an HTH domain.

### Phylogenetic analysis

The tree in Figure 5B holds one intact homolog per cluster. We aligned the full-length sequences with MAFFT 7.526^39^ (L-INS-i), removed columns with fewer than 90% of sequences present with trimAl 1.5.1^40^ (428 columns kept), and inferred a maximum-likelihood tree with IQ-TREE 3.1.3^41^ (LG+F+R6; 1,000 SH-aLRT replicates and 1,000 ultrafast bootstraps). For display, we rooted the tree by minimal ancestor deviation (MAD) and visualized it with TreeViewer^72^; colored clades are the largest below the root with SH-aLRT ≥ 80 and ultrafast bootstrap ≥ 95. Note that, initially, the three experimentally tested homologs (Lac, Cfe and Mfo) sat on long, isolated branches due to what’s known as long branch attraction, and landed next to each other. To anchor them better, we included 142 close relatives from the earlier search and later removed them from the tree before generating Figure 5B.

### Genomic contexts and flanks

For each homolog, we retrieved up to 100 genomic contexts using IPG, each from a different genome assembly, and retained only those whose annotated coding sequence translated exactly to the homolog. We cut 1,000 nucleotides (nt) on each side of the gene, oriented to the gene so that the upstream flank ends at the start codon and the downstream flank begins at the stop codon. We excluded flanks with fewer than 100 usable nt and those with bases other than A, C, G or T. Since closely related strains often carry the same gene in nearly the same neighborhood, and counting every copy would allow a few heavily sequenced species outvote the rest, we merged same-side flanks of a homolog that differed by at most 2% edits using Edlib 1.3.9^71^. When we later pooled the flanks of a cluster, we used the same rule to set aside near-copies coming from different members of that cluster.

### Repeat detection

We called tandem arrays with Tandem Repeats Finder (TRF) 4.09 (match, mismatch and indel weights 2, 7 and 7; match and indel probabilities 80 and 10; minimum score 50; at least two full copies). TRF only detects copies that reside directly next to one another. We adopted Vmatch 2.3.1^43^ (a versatile software tool originally intended for efficiently solving large scale sequence matching tasks) to detect spaced repeats by comparing each flank’s index against itself. Note that while TRF reports all copies of a multi-copy array at once, Vmatch used in this way reports matches in pairs (e.g., a three-copy repeat A-B-C is reported as A-B, B-C and A-C). Since we defined a flank’s repeat detection outcome as whether repeat DNA covers bases within a flank, a spaced repeat counts the same whether it is read as one multi-copy repeat structure, or as the pairs that comprise it, we therefore counted the pairs directly. Note that we also ignored direct pairs from Vmatch lying inside a TRF array, as pieces of that array already detected to avoid double counting.

Within each flank, we searched with Vmatch for direct pairs whose copies lie at most 300 nt apart and for inverted pairs at any distance (we excluded self-palindromes from the analysis), extending exact seeds of 10 nt to at least 80% identity and keeping pairs with a loose Vmatch E-value of 0.01 or less, for later rescoring. To enhance stringency, the Vmatch searches were permissive of mismatches, but not insertions or deletions (i.e., Hamming distance), whereas TRF was allowed to tolerate small insertions and deletions within tandem arrays.

Note that Vmatch scores each pair with an E-value that assumes all four bases are equally common. For spaced candidate pairs as far apart as 300 nt, we were concerned that this would likely underestimate how often repeats arise by chance in DNA of skewed composition. To remedy this, we recomputed each pair’s E-value with a first-order Markov model we derived beyond the simplistic analytical formula used by Vmatch^43^, as described in the next section below. This model not only accounts for how often each base actually occurs but also models base-to-base transition frequencies with a matrix. Based on this E-value model, we kept only pairs with an E-value of 10⁻³ or less, that is, pairs expected by chance at most once in a thousand flanks. To choose this cutoff, we tested values from 10⁻⁸ to 10⁻² and tracked which repeat pairs each iteration removed, sorting flanks by sequence complexity (entropy rate) and AT content, and matched pairs by the DUST low-complexity score of their copies (Morgulis et al., 2006), without reference to which homolog a flank belongs to. Looser cutoffs removed pairs overwhelmingly from simple, AT-rich flanks, where chance matches come easily; and beyond 10⁻³ however, stricter cutoffs removed pairs at the same rate regardless of flank composition. TRF arrays carry no E-value and were counted as called. However, the analytical formula would estimate that a two-copy array (rarely observed in our dataset) is ∼10³x less likely to occur in tandem compared to an (average) spaced arrangement, with the E-value decreasing rapidly as copy number increases further.

### Markov E-value calculation

For each searched flank, we estimated a first-order transition matrix from its dinucleotide counts. If C_ab_ is the number of times base b follows base a, we used:

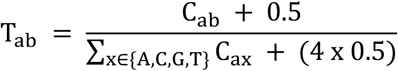

The pseudocount of 0.5 keeps unobserved transitions possible. We obtained the stationary base frequencies π from πT = π, where ∑_x∈{A,C,G,T}_ π_x_ = 1.

For r copies of length *P*, we calculated q*_P_*_,k,r_: the probability that sequences generated independently from this background would resemble one another by chance, with each copy differing from the first at no more than k positions. We also required that the copies could not all be extended by another matching base at either end, so that a shorter piece inside a longer repeat would not be counted separately. These boundary conditions count repeats that cannot be extended by another base shared by all copies. We evaluated this probability using dynamic programming, tracking the current base in each copy and its mismatch count relative to the first.

Forward transitions determine the right boundary; reverse transitions 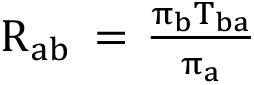 determine the left boundary.

For a searched flank of length n, the approximate (expected) number of such repeats is:

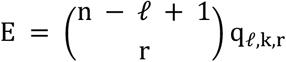

The binomial coefficient counts all unordered sets of r possible copy starts. We used r = 2 to score the repeat pairs reported by Vmatch, with k equal to the actual observed mismatch count. The same score was applied to inverted pairs, using their reverse-complement mismatch count with the same transition matrix.

### Selection of background (randomly-selected) genes

Some genomes simply carry more repeats than others. This variation in repeat abundance among genomes necessitates assessing whether repeats are enriched near StaN homologs relative to other genes within the same genome. As background, we drew twenty random genes from each RefSeq genome holding a homolog flank. We used complete protein-coding genes only, excluding (i) pseudogenes, (ii) genes in several parts, (iii) homologs in this analysis and (iv) genes whose 1-kb flanks overlap those of a homolog. Homologs that their own genome annotates as partial or as a pseudogene were removed by the same rules. In all, 3,234 RefSeq genomes held a homolog flank that entered the comparison, with 64,680 background genes. Background flanks went through the same analysis as homolog flanks. Genomes from the curated RefSeq database were analyzed separately from the 545 genomes found only in GenBank, whose annotations we deemed less reliable. A flank was included in the count for a given distance (e.g., 200, 500, or 1000 nt) only if it had been searched over that entire distance. Within 200 nt, the RefSeq comparison held 2,868 HTH+ flanks (701 clusters, 1,786 genomes, 269 genera) and 2,496 HTH++ flanks (784 clusters, 1,409 genomes, 402 genera) against 125,358 background flanks from the same genomes.

### Repeat analysis by side, distance, size and family

To characterize the spatial distribution of repeats, we repeated the comparison for upstream and downstream flanks separately and in three distance bands, 0–200, 200–500 and 500–1,000 nt (Figure 5G). For repeat size (Figure 5D), the outcome is whether counted repeats cover at least a given length (x axis) within 200 nt, with curves stopping where fewer than 10 clusters reach that length; for unit length (Figure 5H), it is whether a flank carries a tandem array of a given unit length within 200 nt, in 1-nt steps up to 150 nt. For HTH families (Figure 5F), each homolog was counted under every HTH family it carries, we report families with at least 20 clusters in at least 10 genera within a class, and we fitted the Bayesian model separately for each family.

### GC content analysis

AT-rich DNA is a common mark of genes acquired from elsewhere, so we compared each flank’s GC with the mean of background flanks of the same genome and side, adjusted for the distance to the neighboring gene. For Figure 5I, we cut each whole context (1,000 nt upstream, the gene and 1,000 nt downstream) from the genomes and compared the GC of each part, and of the gene’s third codon positions (GC3), with background contexts of the same genome.

### Quantification and statistical analysis of StaN homologs

We used two complementary approaches: a direct, equally weighted comparison across homolog clusters (the “Counting estimate” section below), with uncertainty estimated by resampling clusters and host genera, and a Bayesian model that accounts for genome and cluster differences while stabilizing estimates for sparsely sampled clusters. Agreement between them supports the robustness of our results which show enrichment.

#### Counting estimate

Our approach compares each homolog flank with background flanks from the same genome, accounting for the host’s base composition, genome-wide repeat content and assembly quality. For each comparison, we applied the same repeat-type, location/window and filtering criteria to homolog and background flanks. Let y_0_ ∈ {0,1} indicate whether (homolog) flank i contains qualifying repeat evidence, and b_g(0),4(0)_ be the proportion of eligible background flanks in its genome g(i), on its side d(i), that contain qualifying repeat evidence. For each class (HTH+ or HTH++), we averaged these outcomes and matched background proportions within each cluster, then equally across clusters, to obtain the homolog and background repeat-bearing proportions. With S clusters and n_s_ eligible flanks in cluster s, their difference is 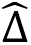 and their ratio is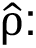

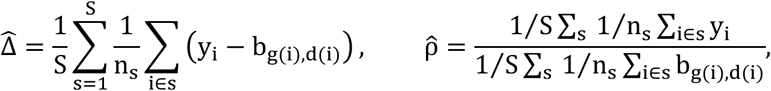

Note that we calculated the reported proportions, differences and enrichment ratios from all eligible flanks. However, although near-identical flanks were collapsed in the previous stages of the pipeline, repeat presence could still be correlated among the remaining flanks of a homolog cluster or host genus. To estimate and report this uncertainty, homolog clusters and host genera were resampled independently, with replacement, 2,000 times. In each resample, we recalculated both proportions, their difference and their ratio, averaging each cluster’s flanks over the genera drawn and weighting clusters by how often they were drawn. The 2.5th and 97.5th percentiles defined the 95% bootstrap intervals. In counting-based plots, estimates come from the full dataset; error bars and the outer stripes of Fig. 5G show the bootstrap interval limits. Two-sided p-values were calculated from how often the resampled homolog-minus-background proportion difference fell on either side of zero, with a minimum reported value of 0.0005. The Bayesian panels (described below) instead show posterior estimates and credible intervals.

#### Bayesian model

We fitted a Bayesian binomial generalized linear mixed model (GLMM) with a complementary log–log (cloglog) link, with random effects for genome and homolog cluster. We organized the flank observations into rows, which we call cells. For homologs, a cell usually contains one flank, identified by its (i) HTH class, (ii) cluster, (iii) host genome and (iv) side. When several homolog flanks share all four labels, they occupy the same cell. For background genes, a cell contains the flanks from one genome and side. Each cell r holds n_r_ flanks, of which k_r_ carry a qualifying repeat. We modeled k_r_ ∼ Binomial(n_r_, p_r_), where p_r_ is the probability that a flank of cell r carries a repeat, and linked p_r_ to the terms of the model through the cloglog link:

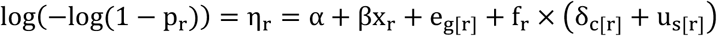

where:

• η is the model’s score for cell r on the cloglog scale; p = 1 − e^−eηr^

• α is the intercept, the score of a background flank in a genome of average repeat content; midway between the two sides, on the model’s scale.

• x_r_ codes the side of cell r (−0.5 upstream, +0.5 downstream), and β is the side effect size;

• g[r] is the genome of cell r, and e_g_ ∼ N(0, σ^2^) is the effect of genome g, how much more or less repeat-prone it is than the average genome, shared by its homolog and background flanks;

• c[r] is the HTH class of cell r (no HTH, HTH+ or HTH++), and δ_c_ is the class shift. It captures how much more or less repeat-prone the flanks of class c are than background flanks of the same genome and side;

• s[r] is the cluster of cell r, and u_s_ ∼ N(0, σ^2^) is the effect of cluster s, its own deviation from its class shift;

• σ_g_ and σ_s_ set how much genomes and clusters vary.

• f_r_ is 1 for homolog cells and 0 for background cells, so the last two terms apply to homolog flanks only;

The cloglog link was chosen since, under a Poisson model, it connects the expected number of events in a flank with the probability of seeing at least one. We also used weakly informative regularizing priors to stabilize estimates where few flanks were available, while allowing substantial differences between groups. For the intercept, we used α ∼ t_3_(−3,2.5), centered on a background repeat probability of about 5%, with a broad spread and heavy tails. For the class shifts and side effect, we used δ_c_, β ∼ N(0,1), centered on no difference but allowing large increases or decreases. For the genome and cluster spreads, we used σ_g_, σ_s_ ∼ Exponential(ln 20/5), favoring smaller differences while giving each spread a 5% prior probability of exceeding 5 on the model’s scale.

The model is ultimately set up to measure the HTH class shift δ_c_ from the flank data and allow us to estimate how much more or less likely the flanks of a given homolog class (HTH+ or HTH++) are to contain qualifying repeats (e.g., tandem, spaced, inverted) than ordinary-gene flanks from the same genomes and sides. We had to address two complications of the data. (1) The number of genomic contexts that were available to download for each homolog ranged from a single genome to many thousands (we set a cap of 100), largely reflecting how often its host species has been sequenced. Even after merging near-identical flanks, clusters still contributed anywhere from one flank to many. A cluster with a single flank gives us little information about its repeat probability, while simply pooling all flanks would give heavily sampled clusters a larger share of the result. (2) Just as the counting estimate approach above, since genomes may differ in how many repeats they carry overall, each homolog needs to be compared with background genes of its own genome.

The key to simultaneously handling these complications was that homolog flanks and background flanks went into one model together. Namely, we described every single flank by its genome, its side, and whether it belongs to a homolog, and if so, to which cluster and class. The model is then able to describe the “probability that a flank carries a repeat” using contributions from different factors: a baseline, its side, its genome, and, for homolog flanks only, a class shift and its cluster’s own deviation. To address the first complication, the cluster term lets each cluster depart from its class’s prediction but draws that departure toward zero when the cluster has few flanks. This helps stabilize estimates for sparsely sampled clusters. Clusters with more flanks provide more information during fitting; and once these cluster estimates have been stabilized using the shared information, we give every cluster equal weight when calculating the reported enrichment, so that heavily sampled clusters do not receive a larger share of the average. To address the second complication, we sampled twenty background genes per genome, each contributing its eligible upstream and downstream flanks. The genome term learns how repeat-rich that genome is, while the class shift and cluster effect together account for how much more or less repeat-prone its homolog flanks are. In this way, each genome serves as its own control.

Since the class shift δ_c_ is added on the model’s transformed scale, log(−log(1 − p)), which keeps every probability between 0 and 1, it is not itself a difference in repeat-bearing proportions. To report it in the same terms as the counting estimate, we interrogated the fitted model with the question: how often would these flanks carry a repeat as homolog flanks, and how often as ordinary-gene flanks of the same genomes and sides? For each homolog cell, p_r_ answers the first one. For the second, we removed the class shift and the cluster effect, η_r_^0^ = η_r_ − f_r_ × (δ_c[r]_ + u_s[r]_), which gives P_r_^0^=1-e-η_r_^0^, the chance for a background gene of the same genome and side. We averaged both over the homolog cells of class c, giving each cell r the weight W_r_ = n_r_/(S n_s[r]_), where S is the number of clusters in the class and n_s[r]_ the number of flanks of cell r’s cluster (background cells and cells of other classes weigh 0). The weights sum to 1, and, as in the counting estimate, flanks count equally within a cluster and clusters count equally. Note that these cell weights give each cluster a total weight of 1/S, divided equally among its eligible flanks. For each posterior draw, we calculated the difference Δ and the enrichment ratio ρ, We report their posterior medians and 95% credible intervals:

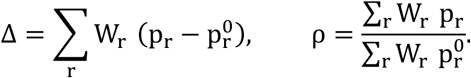

We sampled with CmdStan 2.39.0 through cmdstanr 0.9.0 (R 4.5.3; 4 chains, 1,000 warm-up and 1,000 sampling iterations each) and report posterior medians with 95% credible intervals. As a check using a different fitting approach, we refitted the same model by maximum likelihood with glmmTMB 1.1.14, which gave closely matching estimates (Figure 5E, triangles).

### Lineage cross-check

A repeat beside a homolog could belong to the protein, traveling with it from host to host, or it could simply reflect the host: some genera may be rich in repeats, and any gene living there would have them too. To quantify this, we compared the flanks of each HTH+ homolog across the genomes that carry it, and shuffled repeat-presence labels among HTH+ homolog flanks of the same host genus (2,000 shuffled datasets). This allowed each genus to keep its own repeat rate, so a repeat-rich genus stayed repeat-rich, but any link between a repeat label and a particular HTH+ homolog was broken. The result was that the repeat presence remained associated with the homolog cluster across host genera: for instance, 25% of the HTH+ clusters with four or more distinct upstream flanks had a repeat in all of them, nearly three times the 9% share the shuffling predicted. Furthermore, among pairs of upstream flanks from the same homolog cluster but different host genera, when the first flank carried a repeat, the second carried one too in 92% of pairs, against 43% after shuffling (p = 0.0005 for both comparisons), indicating a near-consistent repeat presence across hosts, well above what we would expect from the hosts’ own tendency to carry repeats.

### Quantification and Statistical Analyses of Experimental Results

Student’s t-tests were used to determine if differences in log transformed plaque counts (Figures 1 & 3 and Supplementary Figures 1 & 7), log transformed colony counts (Figures 2, 4, 6 and Supplementary Figure 7), DNA abundance (Figure 1), Relative Fluorescence Units (Supplementary Figure 7) and enzymatic assays (Figure 3) were statistically significant. Tests were performed on Microsoft excel, and p values below 0.05 were deemed significant. All statistics used for the homolog and repeat analyses in Figure 5 are described in the section above.

### Data and code availability

Raw Illumina sequencing reads for phage ISP and mutant variants have been deposited at NCBI Sequence Read Archive (SRA) under BioProject PRJNA1522937 and are publicly available as of the date of publication. Accession numbers are listed in the key resources table. All other data reported in this paper will be shared by the lead contact upon request. Computational analyses were conducted using publicly-available software packages and cited in the appropriate sections of the narrative. Any additional information required to reanalyze the data reported in this paper is available from lead contact upon request.

## Supplemental Information

The following supplementary files are also available:

**Supplementary Figure 1.** Related to Figure 1. SCC*mec*-encoded gene(s) defend against K-like myophages.

**Supplementary Figure 2.** Related to Figure 1. Three myophages used in this study are similar to staphylococcal phage K.

**Supplementary Figure 3.** Related to Figure 3. Purification schematic and size exclusion chromatograms of StaN and mutant variants.

**Supplementary Figure 4.** Related to Figure 3. tRNA cleavage activity of StaN in the presence of different metals.

**Supplementary Figure 5.** Related to Figure 4. Phage ISP does not degrade host DNA over the course of infection.

**Supplementary Figure 6.** Related to Figure 4. Illumina sequencing reads of 11 ISP escaper phages compared to ancestral phage ISP.

**Supplementary Figure 7.** Related to Figure 4. AlphaFold 3 models of Gp85 and StaN bound to various nucleotide sequences.

**Table S1.** Accompanies Figure 3A. 717 StaN homologs for amino acid conservation analysis

**Table S2.** Accompanies Figure S7 A and B. ISP Promoter elements and AlphaFold 3 confidence scores when the double-stranded DNA sequences are folded with ISP gp_85

**Table S3.** Accompanies Figure S7 C and D. iPTM and pTM scores of ISP Gp85 and StaN bound to 18-nucleotide stretches of the *staI* locus

**Table S4.** Accompanies Figure 5. Summary of StaN homologs for protein architecture and genome context analysis

**Table S5.** Accompanies Figure 5. Summary of flanking regions used in repeat analysis

**Table S6.** Accompanies STAR Methods. DNA oligonucleotides for PCR and Sanger sequencing

**Table S7.** Accompanies STAR Methods. Synthetic constructs.

**Table S8.** Accompanies STAR Methods. DNA oligonucleotides (PAGE purified) for biochemical assays.

## Notes

### Competing Interest Statement

The authors have declared no competing interest.

## References

1. Boyle, T.A., and Hatoum-Aslan, A. (2023). Recurring and emerging themes in prokaryotic innate immunity. Preprint at Elsevier Ltd, 10.1016/j.mib.2023.102324.

2. Rousset, F., and Sorek, R. (2023). The evolutionary success of regulated cell death in bacterial immunity. Preprint at Elsevier Ltd, 10.1016/j.mib.2023.102312.

3. Harms, A., Brodersen, D.E., Mitarai, N., and Gerdes, K. (2018). Toxins, Targets, and Triggers: An Overview of Toxin-Antitoxin Biology. Preprint at Cell Press, 10.1016/j.molcel.2018.01.003.

4. Leroux, M., and Laub, M.T. (2022). Toxin-Antitoxin Systems as Phage Defense Elements. Annu. Rev. Microbiol. 36, 41. 10.1146/annurev-micro-020722.

5. Jurėnas, D., Fraikin, N., Goormaghtigh, F., and Van Melderen, L. (2022). Biology and evolution of bacterial toxin–antitoxin systems. Preprint at Nature Research, 10.1038/s41579-021-00661-1.

6. Wang, X., Lord, D.M., Cheng, H.Y., Osbourne, D.O., Hong, S.H., Sanchez-Torres, V., Quiroga, C., Zheng, K., Herrmann, T., Peti, W., et al. (2012). A new type V toxin-antitoxin system where mRNA for toxin GhoT is cleaved by antitoxin GhoS. Nat. Chem. Biol. 8, 855–861. 10.1038/nchembio.1062.

7. Aakre, C.D., Phung, T.N., Huang, D., and Laub, M.T. (2013). A bacterial toxin inhibits DNA replication elongation through a direct interaction with the β sliding clamp. Mol. Cell 52, 617–628. 10.1016/j.molcel.2013.10.014.

8. Songailiene, I., Juozapaitis, J., Tamulaitiene, G., Ruksenaite, A., Šulčius, S., Sasnauskas, G., Venclovas, Č., and Siksnys, V. (2020). HEPN-MNT Toxin-Antitoxin System: The HEPN Ribonuclease Is Neutralized by OligoAMPylation. Mol. Cell 80, 955–970.e7. 10.1016/j.molcel.2020.11.034.

9. Li, M., Gong, L., Cheng, F., Yu, H., Zhao, D., Wang, R., Wang, T., Zhang, S., Zhou, J., Shmakov, S.A., et al. (2021). Toxin-antitoxin RNA pairs safeguard CRISPR-Cas systems. Science (1979). 372. 10.1126/science.abe5601.

10. Millman, A., Bernheim, A., Stokar-Avihail, A., Fedorenko, T., Voichek, M., Leavitt, A., Oppenheimer-Shaanan, Y., and Sorek, R. (2020). Bacterial Retrons Function In Anti-Phage Defense. Cell 183, 1551–1561.e12. 10.1016/j.cell.2020.09.065.

11. Hossain, M., Aslan, B., and Hatoum-Aslan, A. (2024). Tandem mobilization of anti-phage defenses alongside SCCmec elements in staphylococci. Nature Communications 15. 10.1038/s41467-024-53146-z.

12. Bongiorno, D., Campanile, F., Mongelli, G., Baldi, M.T., Provenzani, R., Reali, S., Lo Russo, C., Santagati, M., and Stefani, S. (2010). DNA methylase modifications and other linezolid resistance mutations in coagulase-negative staphylococci in Italy. Journal of Antimicrobial Chemotherapy 65, 2336–2340. 10.1093/jac/dkq344.

13. Horinouchit, S., and Weisblum, B. (1982). Nucleotide Sequence and Functional Map of pC194, a Plasmid That Specifies Inducible Chloramphenicol Resistance. J. Bacteriol., 815–825.

14. Nair, D., Memmi, G., Hernandez, D., Bard, J., Beaume, M., Gill, S., Francois, P., and Cheung, A.L. (2011). Whole-genome sequencing of Staphylococcus aureus strain RN4220, a key laboratory strain used in virulence research, identifies mutations that affect not only virulence factors but also the fitness of the strain. J. Bacteriol. 193, 2332– 2335. 10.1128/JB.00027-11.

15. Jiang, W., Maniv, I., Arain, F., Wang, Y., Levin, B.R., and Marraffini, L.A. (2013). Dealing with the Evolutionary Downside of CRISPR Immunity: Bacteria and Beneficial Plasmids. PLoS Genet. 9. 10.1371/journal.pgen.1003844.

16. Depardieu, F., Didier, J.P., Bernheim, A., Sherlock, A., Molina, H., Duclos, B., and Bikard, D. (2016). A Eukaryotic-like Serine/Threonine Kinase Protects Staphylococci against Phages. Cell Host Microbe 20, 471–481. 10.1016/j.chom.2016.08.010.

17. Bari, S.M.N., Chou-Zheng, L., Howell, O., Hossain, M., Hill, C.M., Boyle, T.A., Cater, K., Dandu, V.S., Thomas, A., Aslan, B., et al. (2022). A unique mode of nucleic acid immunity performed by a multifunctional bacterial enzyme. Cell Host Microbe 30, 570–582.e7. 10.1016/j.chom.2022.03.001.

18. Barylski, J., Kropinski, A.M., Alikhan, N.F., and Adriaenssens, E.M. (2020). ICTV virus taxonomy profile: Herelleviridae. Journal of General Virology 101, 362–363. 10.1099/jgv.0.001392.

19. Łobocka, M., Hejnowicz, M.S., Dabrowski, K., Gozdek, A., Kosakowski, J., Witkowska, M., Ulatowska, M.I., Weber-Dabrowska, B., Kwiatek, M., Parasion, S., et al. (2012). Genomics of Staphylococcal Twort-like Phages-Potential Therapeutics of the Post-Antibiotic Era. In Advances in Virus Research (Academic Press Inc.), pp. 143–216. 10.1016/B978-0-12-394438-2.00005-0.

20. Carroll, R.K., Weiss, A., Broach, W.H., Wiemels, R.E., Mogen, A.B., Rice, K.C., and Shaw, L.N. (2016). Genome-wide annotation, identification, and global transcriptomic analysis of regulatory or small RNA gene expression in Staphylococcus aureus. mBio 7. 10.1128/mBio.01990-15.

21. Fineran, P.C., Blower, T.R., Foulds, I.J., Humphreys, D.P., Lilley, K.S., and Salmond, G.P.C. (2009). The phage abortive infection system, ToxIN, functions as a protein-RNA toxin-antitoxin pair. Proc. Natl. Acad. Sci. U. S. A. 106, 894–899.

22. Blower, T.R., Short, F.L., Rao, F., Mizuguchi, K., Pei, X.Y., Fineran, P.C., Luisi, B.F., and Salmond, G.P.C. (2012). Identification and classification of bacterial Type III toxin-antitoxin systems encoded in chromosomal and plasmid genomes. Nucleic Acids Res. 40, 6158–6173. 10.1093/nar/gks231.

23. Goeders, N., Chai, R., Chen, B., Day, A., and Salmond, G.P.C. (2016). Structure, evolution, and functions of bacterial type III toxin-antitoxin systems. Toxins (Basel). 8. 10.3390/toxins8100282.

24. Naville, M., Ghuillot-Gaudeffroy, A., Marchais, A., and Gautheret, D. (2011). ARNold: A web tool for the prediction of rho-independent transcription terminators. RNA Biol. 8, 11–13. 10.4161/rna.8.1.13346.

25. Hatoum-Aslan, A., Maniv, I., and Marraffini, L.A. (2011). Mature clustered, regularly interspaced, short palindromic repeats RNA (crRNA) length is measured by a ruler mechanism anchored at the precursor processing site. Proc. Natl. Acad. Sci. U. S. A. 108, 21218–21222. 10.1073/pnas.1112832108.

26. Hatoum-Aslan, A., Samai, P., Maniv, I., Jiang, W., and Marraffini, L.A. (2013). A ruler protein in a complex for antiviral defense determines the length of small interfering CRISPR RNAs. Journal of Biological Chemistry 288, 27888–27897. 10.1074/jbc.M113.499244.

27. Blum, M., Chang, H.Y., Chuguransky, S., Grego, T., Kandasaamy, S., Mitchell, A., Nuka, G., Paysan-Lafosse, T., Qureshi, M., Raj, S., et al. (2021). The InterPro protein families and domains database: 20 years on. Nucleic Acids Res. 49, D344–D354. 10.1093/nar/gkaa977.

28. Perez Taboada, V., Wu, Y., Cassidy, R., Medvedev, K.E., Loeff, L., Nemudraia, A., and Nemudryi, A. (2026). Bacterial Schlafen proteins mediate phage defence. Nat. Microbiol. 11, 1037–1048. 10.1038/s41564-026-02277-8.

29. Weickert, P., Liu, Y., and Strecker, J. (2026). Prokaryotic Schlafen proteins cleave tRNAs during type III CRISPR immunity. Nature Communications 17, 1–12. 10.1038/s41467-026-74880-6.

30. Thomas, P.D., Ebert, D., Muruganujan, A., Mushayahama, T., Albou, L.P., and Mi, H. (2022). PANTHER: Making genome-scale phylogenetics accessible to all. Preprint at John Wiley and Sons Inc, 10.1002/pro.4218.

31. Abramson, J., Adler, J., Dunger, J., Evans, R., Green, T., Pritzel, A., Ronneberger, O., Willmore, L., Ballard, A.J., Bambrick, J., et al. (2024). Accurate structure prediction of biomolecular interactions with AlphaFold 3. Nature 630, 493–500. 10.1038/s41586-024-07487-w.

32. Rees, P.J., and Fry, B.A. (1981). The Morphology of Staphylococcal Bacteriophage K and DNA Metabolism in Infected Staphylococcus aureus. Journal of General Virology 53, 293–307.

33. Finstrlová, A., Mašlaňová, I., Blasdel Reuter, B.G., Doškař, J., Götz, F., and Pantůček, R. (2022). Global Transcriptomic Analysis of Bacteriophage-Host Interactions between a Kayvirus Therapeutic Phage and Staphylococcus aureus. Microbiol. Spectr. 10, 1–13. 10.1128/spectrum.00123-22.

34. Lehman, S.M., Kongari, R., Glass, A.M., Koert, M., Ray, M.D., Plaut, R.D., and Stibitz, S. (2023). Phage K gp102 Drives Temperature-Sensitive Antibacterial Activity on USA300 MRSA. Viruses 15. 10.3390/v15010017.

35. Kongari, R., Ray, M.D., Lehman, S.M., Plaut, R.D., Hinton, D.M., and Stibitz, S. (2024). The Transcriptional Program of Staphylococcus aureus Phage K Is Affected by a Host rpoC Mutation That Confers Phage K Resistance. Viruses 16. 10.3390/v16111773.

36. Berger, M.F., Badis, G., Gehrke, A.R., Talukder, S., Philippakis, A.A., Peña-Castillo, L., Alleyne, T.M., Mnaimneh, S., Botvinnik, O.B., Chan, E.T., et al. (2008). Variation in Homeodomain DNA Binding Revealed by High-Resolution Analysis of Sequence Preferences. Cell 133, 1266–1276. 10.1016/j.cell.2008.05.024.

37. Zhang, J., Schaeffer, R.D., Durham, J., Cong, Q., and Grishin, N. V. (2023). DPAM: A domain parser for AlphaFold models. Protein Science 32. 10.1002/pro.4548.

38. Cheng, H., Schaeffer, R.D., Liao, Y., Kinch, L.N., Pei, J., Shi, S., Kim, B.H., and Grishin, N. V. (2014). ECOD: An Evolutionary Classification of Protein Domains. PLoS Comput. Biol. 10. 10.1371/journal.pcbi.1003926.

39. Katoh, K., and Standley, D.M. (2013). MAFFT multiple sequence alignment software version 7: Improvements in performance and usability. Mol. Biol. Evol. 30, 772–780. 10.1093/molbev/mst010.

40. Capella-Gutiérrez, S., Silla-Martínez, J.M., and Gabaldón, T. (2009). trimAl: A tool for automated alignment trimming in large-scale phylogenetic analyses. Bioinformatics 25, 1972–1973. 10.1093/bioinformatics/btp348.

41. Wong, T.K.F., Ly-Trong, N., Ren, H., Demotte, P., Baños, H., Roger, A.J., Susko, E., Bielow, C., De Maio, N., Goldman, N., et al. (2026). IQ-TREE 3: phylogenomic inference software using complex evolutionary models. Mol. Biol. Evol. 43. 10.1093/molbev/msag117.

42. Benson, G. (1999). Tandem repeats finder: a program to analyze DNA sequences. Nucleic Acids Res. 27, 573–580.

43. Fonseca, N.A., Rung, J., Brazma, A., and Marioni, J.C. (2012). Tools for mapping high-throughput sequencing data. Preprint, 10.1093/bioinformatics/bts605.

44. Muto, A., and Osawa, S. (1987). The guanine and cytosine content of genomic DNA and bacterial evolution (biased mutation pressure/codon usage/neutral theory). Proc. Nati. Acad. Sci. USA 84, 166–169.

45. Knight, R.D., Freeland, S.J., and Landweber, L.F. (2001). A simple model based on mutation and selection explains trends in codon and amino-acid usage and GC composition within and across genomes. Genome Biol. 2.

46. Rocha, E.P.C., and Danchin, A. (2002). Base composition bias might result from competition for metabolic resources. Trends in Genetics 18, 291–294.

47. Kim, E.T., and Weitzman, M.D. (2022). Schlafens Can Put Viruses to Sleep. Viruses 14, 1–20. 10.3390/v14020442.

48. Li, M., Kao, E., Malone, D., Gao, X., Wang, J.Y.J., and David, M. (2018). DNA damage-induced cell death relies on SLFN11-dependent cleavage of distinct type II tRNAs. Nat. Struct. Mol. Biol. 25, 1047–1058. 10.1038/s41594-018-0142-5.

49. Yang, J.-Y., Deng, X.-Y., Li, Y.-S., Ma, X.-C., Feng, J.-X., Yu, B., Chen, Y., Luo, Y.-L., Wang, X., Chen, M.-L., et al. (2018). Structure of Schlafen13 reveals a new class of tRNA/rRNA-targeting RNase engaged in translational control. Nat. Commun. 9, 1–13. 10.1038/s41467-018-03544-x.

50. Seong, R.-K., Seo, S., Kim, J.-A., Fletcher, S.J., Morgan, N. V., Kumar, M., Choi, Y.-K., and Shin, O.S. (2017). Schlafen 14 (SLFN14) is a novel antiviral factor involved in the control of viral replication. Immunobiology 222, 979–988. 10.1016/j.imbio.2017.07.002.

51. Pisareva, V.P., Muslimov, I.A., Tcherepanov, A., and Pisarev, A. V. (2015). Characterization of novel ribosome-associated endoribonuclease SLFN14 from rabbit reticulocytes. Biochemistry 54, 3286–3301. 10.1021/acs.biochem.5b00302.

52. Li, M., Kao, E., Gao, X., Sandig, H., Limmer, K., Pavon-Eternod, M., Jones, T.E., Landry, S., Pan, T., Weitzman, M.D., et al. (2012). Codon-usage-based inhibition of HIV protein synthesis by human schlafen 11. Nature 491, 125–128. 10.1038/nature11433.

53. Bennett, B.D., Kimball, E.H., Gao, M., Osterhout, R., Van Dien, S.J., and Rabinowitz, J.D. (2009). Absolute metabolite concentrations and implied enzyme active site occupancy in Escherichia coli. Nat. Chem. Biol. 5, 593–599. 10.1038/nchembio.186.

54. Deng, Y., Beahm, D.R., Ionov, S., and Sarpeshkar, R. (2021). Measuring and modeling energy and power consumption in living microbial cells with a synthetic ATP reporter. BMC Biol. 19, 1–21. 10.1186/s12915-021-01023-2.

55. Diep, B.A., Gill, S.R., Chang, R.F., Phan, T.H. Van, Chen, J.H., Davidson, M.G., Lin, F., Lin, J., Carleton, H.A., Mongodin, E.F., et al. (2006). Complete genome sequence of USA300, an epidemic clone of community-acquired meticillin-resistant Staphylococcus aureus. Lancet 367, 731–739. 10.1016/S0140-6736(06)68231-7.

56. Vandersteegen, K., Mattheus, W., Ceyssens, P.J., Bilocq, F., de Vos, D., Pirnay, J.P., Noben, J.P., Merabishvili, M., Lipinska, U., Hermans, K., et al. (2011). Microbiological and molecular assessment of bacteriophage ISP for the control of Staphylococcus aureus. PLoS One 6. 10.1371/journal.pone.0024418.

57. Antoine Marc, A., Cater, K., Kongari, R., Hatoum-Aslan, A., III, R.F.Y., and Liu, M. (2019). Complete Genome Sequence of Staphylococcus aureus Siphophage Lorac. Microbiol. Resour. Announc. 8, e00586–19.

58. Culbertson, E.K., Bari, S.M.N., Dandu, S., Kriznik, J.M., Scopel, S.E., Stanley, S.P., Lackey, K., Hernandez, A.C., and Hatoum-Aslan, A. (2019). Draft Genome Sequences of Staphylococcus Podophages JBug18, Pike, Pontiff, and Pabna. Microbiol. Resour. Announc. 8, e00054–19.

59. Freeman, M.E., Kenny, S.E., Lanier, A., Cater, K., Wilhite, M.C., Gamble, P., O’Leary, C.J., Hatoum-Aslan, A., Young, R.F.I., and Liu, M. (2019). Complete Genome Sequences of Staphylococcus epidermidis Myophages Quidividi, Terranova, and Twillingate. Microbiol. Resour. Announc. 8, e00598–19.

60. Hill, C.M., Pollitz, L.J., Boyle, T.A., Akrami, A., Briley, C., Cichanski, S., DeMarco, A., Gonzalez, K., Rojas Nava, N., Qavi, Y., et al. (2026). Complete genome sequences of Staphylococcus epidermidis phages Cicami, Sazerac, Southeast, Slasher, Spartan, and Undine. Microbiol. Resour. Announc. 15. 10.1128/mra.01000-25.

61. Cater, K., Dandu, V.S., Bari, S.M.N., Lackey, K., Everett, G.F.K., and Hatoum-Aslan, A. (2017). A Novel Staphylococcus Podophage Encodes a Unique Lysin with Unusual Modular Design. 2, e00040–17.

62. Charpentier, E., Anton, A.I., Barry, P., Alfonso, B., Fang, Y., and Novick, R.P. (2004). Novel cassette-based shuttle vector system for gram-positive bacteria. Appl. Environ. Microbiol. 70, 6076–6085. 10.1128/AEM.70.10.6076-6085.2004.

63. Bae, T., and Schneewind, O. (2006). Allelic replacement in Staphylococcus aureus with inducible counter-selection. Plasmid 55, 58–63. 10.1016/j.plasmid.2005.05.005.

64. Weisblum, B., Graham, M.Y., Gryczan, T., and Dubnau, D. (1979). Plasmid Copy Number Control: Isolation and Characterization of High-Copy-Number Mutants of Plasmid pE194. J. Bacteriol., 635–643.

65. Samai, P., Pyenson, N., Jiang, W., Goldberg, G.W., Hatoum-Aslan, A., and Marraffini, L.A. (2015). Co-transcriptional DNA and RNA cleavage during type III CRISPR-cas immunity. Cell 161, 1164–1174. 10.1016/j.cell.2015.04.027.

66. Guzman, L.M., Belin, D., Carson, M.J., and Beckwith, J. (1995). Tight regulation, modulation, and high-level expression by vectors containing the arabinose PBAD promoter. J. Bacteriol. 177, 4121–4130. 10.1128/jb.177.14.4121-4130.1995.

67. Norrander, J., Kempe, T., and Messing, J. (1983). Construction of improved Ml3 vectors using oligodeoxynucleotide-directed mutagenesis. Gene 26, 101–106.

68. Camacho, C., Coulouris, G., Avagyan, V., Ma, N., Papadopoulos, J., Bealer, K., and Madden, T.L. (2009). BLAST+: Architecture and applications. BMC Bioinformatics 10. 10.1186/1471-2105-10-421.

69. Jones, P., Binns, D., Chang, H.Y., Fraser, M., Li, W., McAnulla, C., McWilliam, H., Maslen, J., Mitchell, A., Nuka, G., et al. (2014). InterProScan 5: Genome-scale protein function classification. Bioinformatics 30, 1236–1240. 10.1093/bioinformatics/btu031.

70. Steinegger, M., and Söding, J. (2016). MMseqs2: sensitive protein sequence searching for the analysis of massive data sets. Preprint, 10.1101/079681.

71. Šošić, M., and Šikić, M. (2017). Edlib: A C/C ++ library for fast, exact sequence alignment using edit distance. Bioinformatics 33, 1394–1395. 10.1093/bioinformatics/btw753.

72. Bianchini, G., and Sánchez-Baracaldo, P. (2024). TreeViewer: Flexible, modular software to visualise and manipulate phylogenetic trees. Ecol. Evol. 14, 1–16. 10.1002/ece3.10873.

73. Carpenter, B., Gelman, A., Hoffman, M.D., Lee, D., Goodrich, B., Betancourt, M., Brubaker, M.A., Guo, J., Li, P., and Riddell, A. (2017). Stan: A probabilistic programming language. J. Stat. Softw. 76, 1–32. 10.18637/jss.v076.i01.

74. Meng, E.C., Goddard, T.D., Pettersen, E.F., Couch, G.S., Pearson, Z.J., Morris, J.H., and Ferrin, T.E. (2023). UCSF ChimeraX: Tools for structure building and analysis. Protein Science 32. 10.1002/pro.4792.

75. Schneider, C.A., Rasband, W.S., and Eliceiri, K.W. (2012). NIH Image to ImageJ: 25 years of image analysis. Nat. Methods 9, 671–675. 10.1038/nmeth.2089.

76. Darling, A.C.E., Mau, B., Blattner, F.R., and Perna, N.T. (2004). Mauve: Multiple alignment of conserved genomic sequence with rearrangements. Genome Res. 14, 1394–1403. 10.1101/gr.2289704.

77. Gilchrist, C.L.M., and Chooi, Y.H. (2021). Clinker & clustermap.js: Automatic generation of gene cluster comparison figures. Bioinformatics 37, 2473–2475. 10.1093/bioinformatics/btab007.

78. Bankevich, A., Nurk, S., Antipov, D., Gurevich, A.A., Dvorkin, M., Kulikov, A.S., Lesin, V.M., Nikolenko, S.I., Pham, S., Prjibelski, A.D., et al. (2012). SPAdes: A new genome assembly algorithm and its applications to single-cell sequencing. Journal of Computational Biology 19, 455–477. 10.1089/cmb.2012.0021.

79. Wick, R.R., Schultz, M.B., Zobel, J., and Holt, K.E. (2015). Bandage: Interactive visualization of de novo genome assemblies. Bioinformatics 31, 3350–3352. 10.1093/bioinformatics/btv383.

80. Li, H., Handsaker, B., Wysoker, A., Fennell, T., Ruan, J., Homer, N., Marth, G., Abecasis, G., and Durbin, R. (2009). The Sequence Alignment/Map format and SAMtools. Bioinformatics 25, 2078–2079. 10.1093/bioinformatics/btp352.

81. Robinson, J.T., Thorvaldsdóttir, H., Winckler, W., Guttman, M., Lander, E.S., Getz, G., and Mesirov, J.P. (2011). Integrative genomics viewer. Nat. Biotechnol. 29, 24–26. 10.1038/nbt0111-24.

82. Christensen, G.D., Baddour, L.M., and Simpson, W.A. (1987). Phenotypic variation of Staphylococcus epidermidis slime production in vitro and in vivo. Infect. Immun. 55, 2870–2877.

83. Gibson, D.G., Young, L., Chuang, R.Y., Venter, J.C., Hutchison, C.A., and Smith, H.O. (2009). Enzymatic assembly of DNA molecules up to several hundred kilobases. Nat. Methods 6, 343–345. 10.1038/nmeth.1318.

84. Monk, I.R., Shah, I.M., Xu, M., Tan, M.W., and Foster, T.J. (2012). Transforming the untransformable: Application of direct transformation to manipulate genetically Staphylococcus aureus and Staphylococcus epidermidis. mBio 3. 10.1128/mBio.00277-11.

85. Chou-Zheng, L., and Hatoum-Aslan, A. (2019). A type III-A CRISPR-Cas system employs degradosome nucleases to ensure robust immunity. Elife, 8:e45393.

86. Ho, S.N., Hunt, H.D., Horton, R.M., Pullen, J.K., and Pease, L.R. (1989). Site-directed mutagenesis by overlap extension using the polymerase chain reaction. Gene 77, 51–59.

87. Cronan, J.E. (2006). A family of arabinose-inducible Escherichia coli expression vectors having pBR322 copy control. Plasmid 55, 152–157. 10.1016/j.plasmid.2005.07.001.

88. Lin-Chao, S., and Bremer, H. (1986). Effect of the bacterial growth rate on replication control of plasmid pBR322 in Escherichia coli. Molecular & General Genetics 203, 143– 149.

89. Miki, T., Yasukochi, T., Nagatani, H., Furuno, M., Orita, T., Yamada, H., Imoto, T., and Horiuchi, T. (1987). Construction of a plasmid vector for the regulatable high level expression of eukaryotic genes in *Escherichia coli*: an application to overproduction of chicken lysozyme. Protein Engineering, Design and Selection 1, 327–332. 10.1093/protein/1.4.327.

90. Langmead, B., and Salzberg, S.L. (2012). Fast gapped-read alignment with Bowtie 2. Nat. Methods 9, 357–359. 10.1038/nmeth.1923.

