## Supplementary Figures 1-7 for "Genomic DNA repeats act as an antitoxin for a widespread Schlafen immune effector"

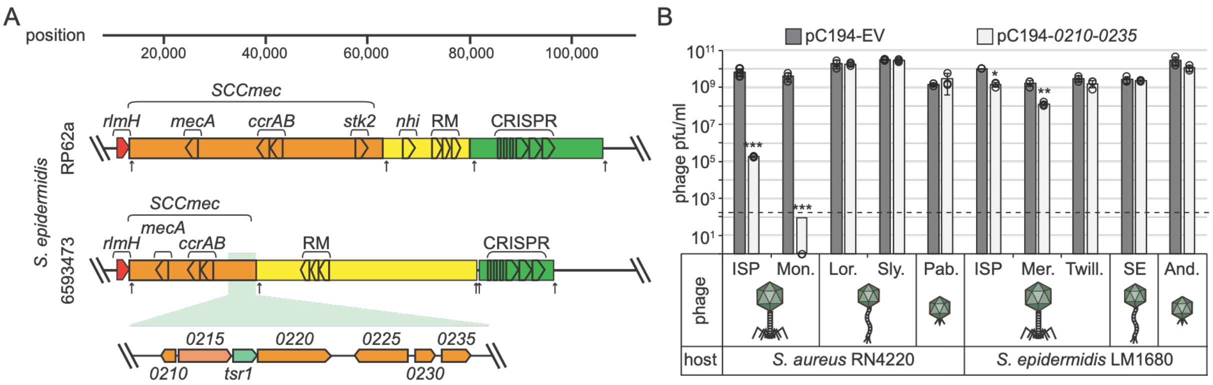


**Figure S1.** Related to Figure 1. SCC*mec*-encoded gene(s) defend against K-like myophages.

**(A)** Diagrams of Staphylococcal Cassette Chromosome (SCC) *mec* and adjacent regions in *S. epidermidis* RP62A and 6593473 genomes. A magnified view of the segment from 6593473 responsible for defense is shown. Arrows indicate CcrAB attachment sites.

**(B)** *S. epidermidis* and *S. aureus* strains bearing a plasmid with the antiphage defense region were challenged with a panel of phages, and the average pfu/ml from three technical replicates are shown (+/- SD).

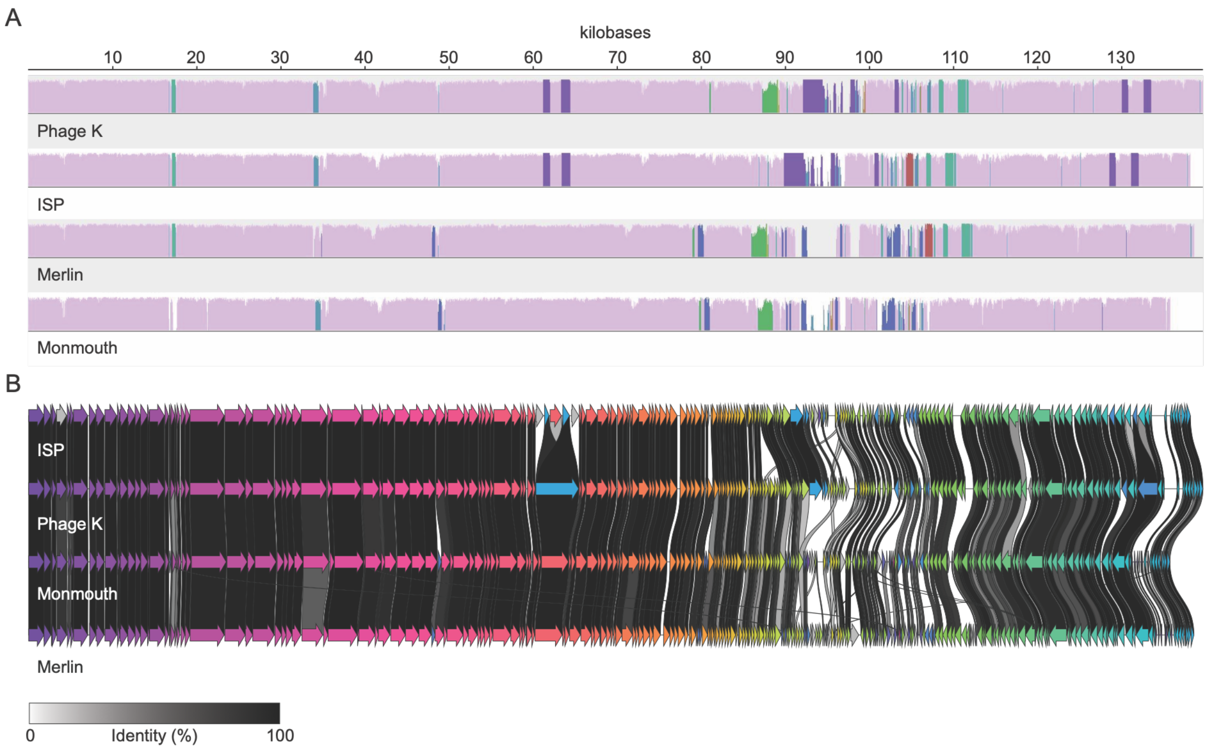


**Figure S2.** Related to Figure 1. Three myophages used in this study are similar to staphylococcal phage K.

**(A)** Mauve (https://darlinglab.org/mauve/mauve.html) and **(B)** Clinker (https://cagecat.bioinformatics.nl/tools/clinker) alignments of phages K, ISP, Merlin, and Monmouth.

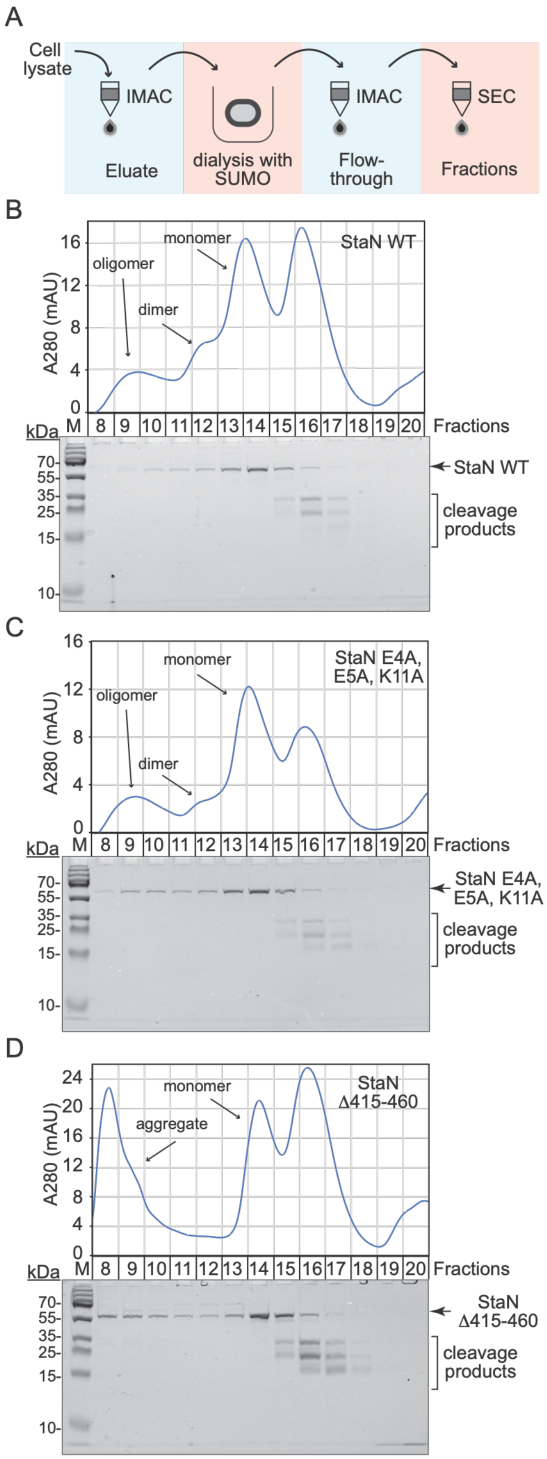
A
Cell lysate
IMAC
Eluate
dialysis with SUMO
IMAC
Flow-through
SEC
Fractions
B
A280 (mAU)
oligomer
dimer
monomer
StaN WT
Fractions
kDa
M
StaN WT
cleavage products
C
StaN E4A, E5A, K11A
oligomer
dimer
monomer
Fractions
StaN E4A, E5A, K11A
cleavage products
D
StaN Δ415-460
monomer
aggregate
Fractions
StaN Δ415-460
cleavage products

**Figure S3.** Related to Figure 3. Purification schematic and size exclusion chromatograms of StaN and mutant variants.

**(A)** Protein purification schematic from *E. coli*. IMAC, immobilized metal affinity chromatography; SUMO, SUMO protease; and SEC, size-exclusion chromatography.

**(B)** – **(D)** Size-exclusion chromatograms of WT StaN and two mutant variants with corresponding SDS-PAGE gels.

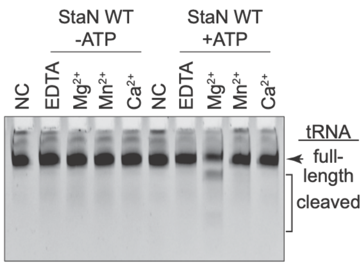


**Figure S4.** Related to Figure 3. tRNA cleavage activity of StaN in the presence of different metals.

tRNA cleavage assays are shown in which tRNA was mixed with WT StaN with or without ATP and different metals, incubated for 40 minutes, and resolved on a Urea-PAGE gel. One representative of three independent trials is displayed.

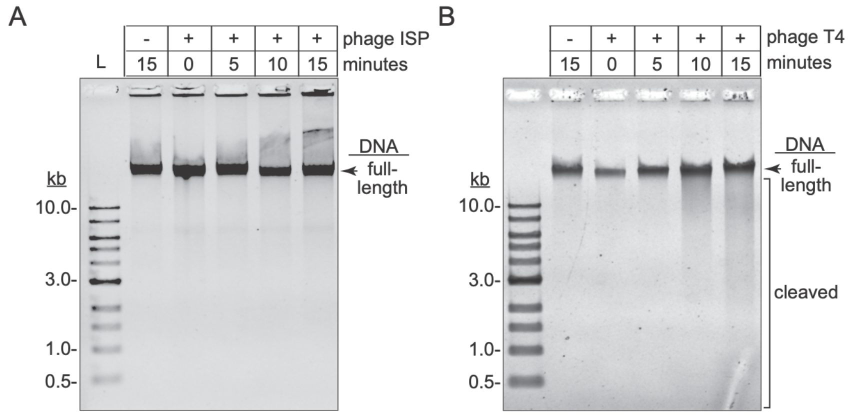


**Figure S5.** Related to Figure 4. Phage ISP does not degrade host DNA over the course of infection.

Results from DNA cleavage assays with phage ISP infection **(A)** and phage T4 infection **(B)**. Total DNA was extracted at 0, 5, 10, and 15 minutes post-infection, resolved on a 0.6% agarose gel, and stained with SYBR™ Gold. One representative assay of three independent trials is displayed.

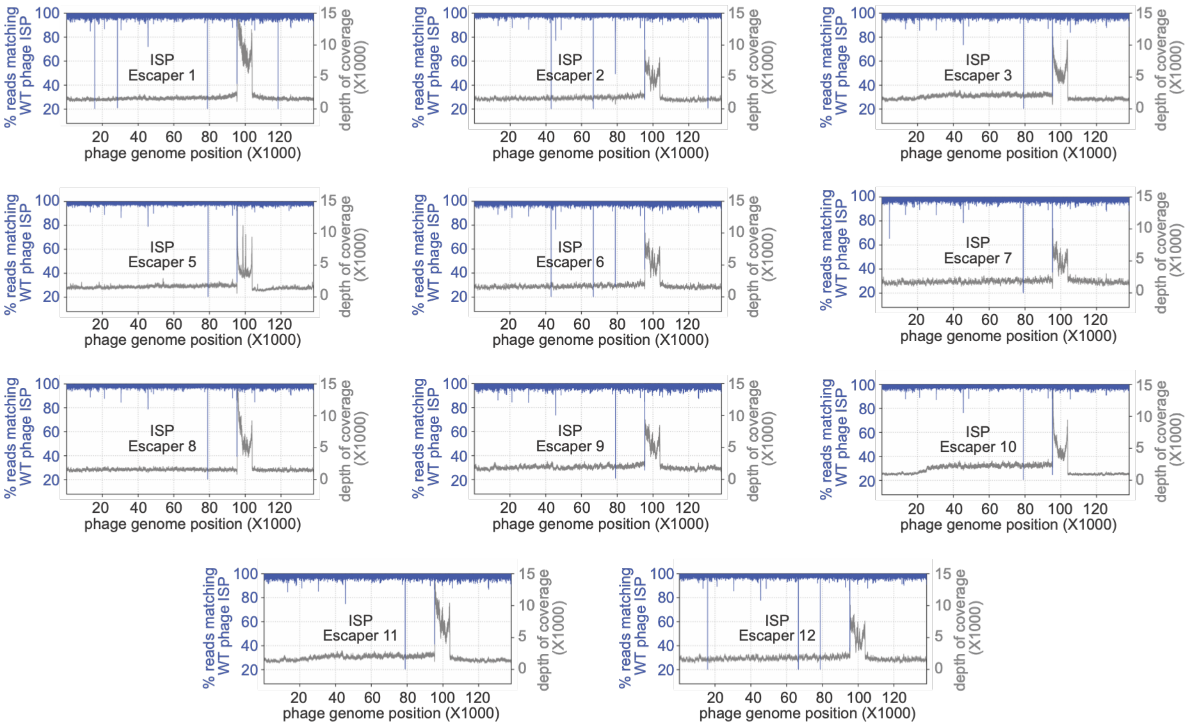


**Figure S6.** Related to Figure 4. Illumina sequencing reads of 11 ISP escaper phages compared to ancestral phage ISP.

Illumina sequencing reads from 11 additional escaper phages aligned with the *de novo* assembly of ancestral ISP. The fraction of reads matching the ancestral ISP was plotted versus the phage genome position.

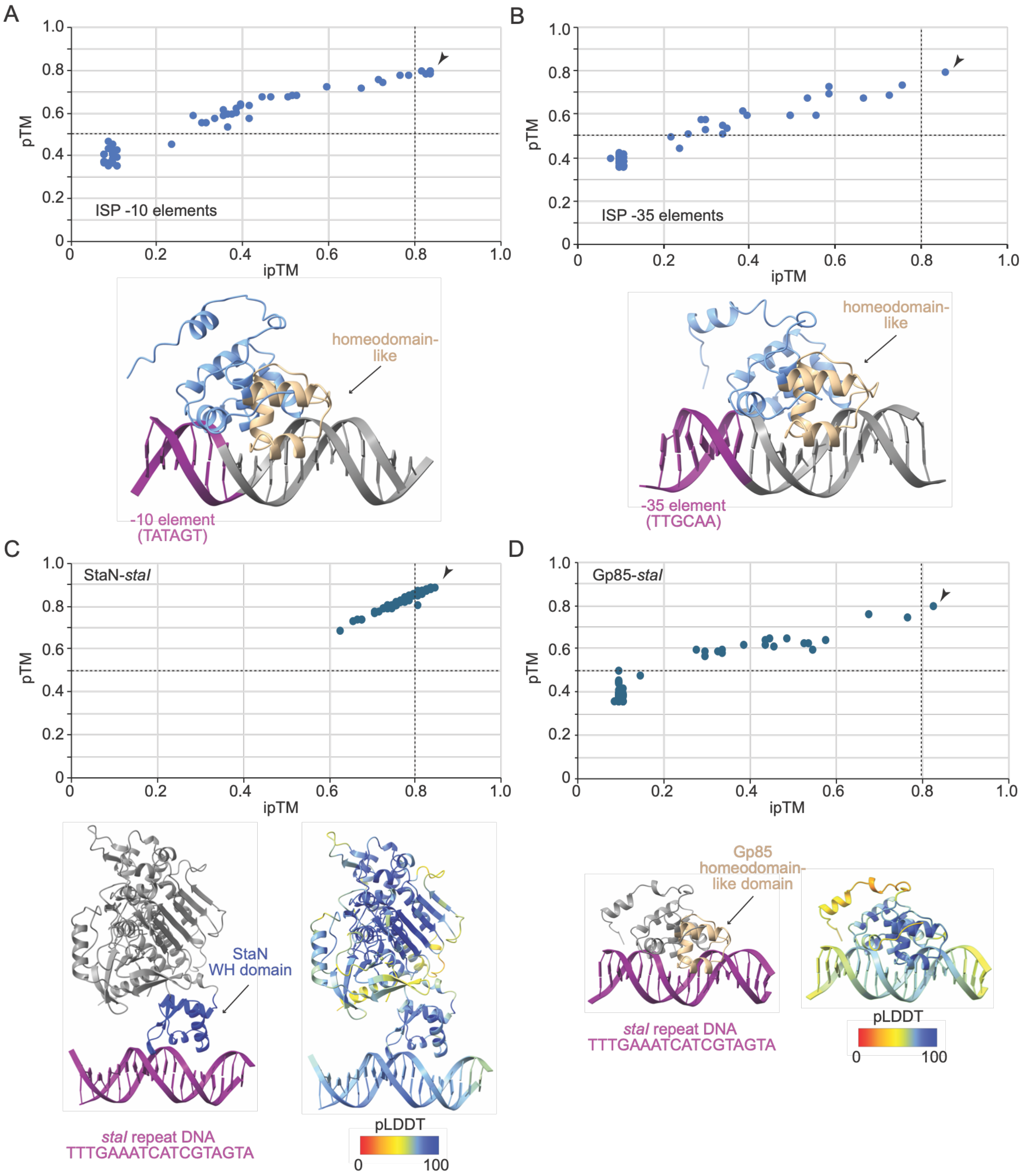
A
B
C
D
pTM
ipTM
ISP -10 elements
ISP -35 elements
homeodomain-like
-10 element (TATAGT)
-35 element (TTGCAA)
StaN-*staI*
Gp85-*staI*
StaN WH domain
*staI* repeat DNA
TTTGAAATCATCGTAGTA
pLDDT
0
100
Gp85 homeodomain-like domain

**Figure S7.** Related to Figure 4. AlphaFold 3 models of Gp85 and StaN bound to various nucleotide sequences.

**(A)** and **(B)** AF3 scan with Gp85 and the -10 and -35 elements of every annotated promoter in the ISP genome.

**(C)** and **(D)** AF3 scan with StaN and Gp85 proteins bound to 18-nt overlapping segments of *staI*.

Predicted template modeling (pTM) scores are plotted versus interface-predicted Template Modeling (ipTM) scores, and the dotted lines indicate the threshold for high-quality, confident predictions of the individual structure or interface. Arrows indicate models with highest confidence that are shown under each corresponding plot. Models are color-coded according to their DNA-binding domains (left) or pLDDT scores (right).
